# QSM Reconstruction Challenge 2.0: Design and Report of Results

**DOI:** 10.1101/2020.11.25.397695

**Authors:** Berkin Bilgic, Christian Langkammer, José P. Marques, Jakob Meineke, Carlos Milovic, Ferdinand Schweser

## Abstract

**Purpose:** The aim of the second quantitative susceptibility mapping (QSM) reconstruction challenge (Oct 2019, Seoul, Korea) was to test the accuracy of QSM dipole inversion algorithms in simulated brain data.

**Methods:** A two-stage design was chosen for this challenge. The participants were provided with datasets of multi-echo gradient echo images synthesized from two realistic *in silico* head phantoms using an MR simulator. At the first stage, participants optimized QSM reconstructions without ground-truths available to mimic the clinical setting. At the second stage, ground-truths were provided for parameter optimization.Submissions were evaluated using eight numerical metrics and visual ratings.

**Results:** A total of 98 reconstructions were submitted for stage 1 and 47 submissions for stage 2. Iterative methods had the best quantitative metric scores, followed by deep-learning and direct inversion methods. Priors derived from magnitude data improved the metric scores. Algorithms based on iterative approaches and Total Variation (and its derivatives) produced the best overall results. The reported results and analysis pipelines have been made public to allow researchers to compare new methods to the current state of the art.

**Conclusion:** The synthetic data provides a consistent framework to test the accuracy and robustness of QSM algorithms in the presence of noise, calcifications and minor voxel dephasing effects. Total Variation-based algorithmsproduced the best results along all metrics. Future QSM challenges should asses if this good performance with synthetic datasets translates to more realistic scenarios, where background fields and dipole-incompatible phase contributions are included.

## Introduction

Quantitative Susceptibility Mapping (QSM) is an emerging MRI technique^1^ that allows for non-invasive estimation of alterations in tissue iron concentration^2,3^, blood oxygenation^4^, and differentiation of paramagnetic and diamagnetic lesions^5,6^. QSM entails the solution of an ill-posed, ill-conditioned inverse problem that relates the acquired gradient echo (GRE) phase information, reflecting magnetic field inhomogeneities, to the underlying susceptibility distribution that is the cause of the inhomogeneities^7,8^. The QSM community has been active in the development of a wide range of reconstruction algorithms^9,10^. These developments may be categorized as inverse filtering (direct/k-space inversion), image-space (regularized iterative reconstruction), and Deep Learning (DL) based approaches. The 2016 QSM Reconstruction Challenge (RC1) provided a common dataset where these algorithms could be compared^11^. This first challenge was highly successful, reflected by 27 submissions from 13 research groups. RC1 used a multi-orientation in vivo dataset This dataset lent itself to susceptibility tensor imaging (STI)^12^ and Calculation Of Susceptibility through Multiple Orientation Sampling (COSMOS) reconstructions^13^, with the goal of mitigating susceptibility anisotropy effects while providing an adequate ground-truth.

RC1 dataset and reconstruction code remained available after the challenge deadline (http://qsm.neuroimaging.at), and found widespread use for benchmarking of new dipole inversion algorithms developed after RC1 had ended^14,15^. Despite its success as a benchmark dataset, the challenge itself had limitations that limited its practical relevance. The limitations were discussed in the report paper, and analyzed more quantitatively in a separate manuscript^16^. In brief, it was concluded that the estimated susceptibility tensor component *χ*_33_ that was used as the ground-truth removed anisotropic contributions found in single-orientation phase data, which resulted in an inconsistency between the provided field map and the groundtruth susceptibility. While the COSMOS solution had higher consistency, the discrepancy was still relatively large and could not be explained just by the noise and background field remnants. Incorporating the *χ_13_* and *χ_23_* anisotropic contributions onto *χ*_33_ would mitigate, but not eliminate this discrepancy. Finding an in vivo ground-truth susceptibility map that matches acquired single-orientation phase data with high fidelity remains as an open problem, further complicated by the presence of background field remnants and the low SNR of the highly accelerated acquisitions. Absence of reliable ground-truth data obstructs quantitative evaluation of the submitted QSM reconstructions.

Our motivation in designing a new reconstruction challenge was the following: (i) understanding the state-of-the-art of QSM algorithms (including the advent of DL techniques^14,15,17–21^ since RC1), plus the identification of limitations of existing algorithms. (ii) objective comparison of published algorithms, incorporating the lessons learnt from RC1, and (iii) providing a new dataset for the future evaluation of dipole inversion algorithms. With these overarching goals, the design of the new challenge, RC2, started during the annual ISMRM meeting in 2018 with a call for ideas. Based on a community driven process, RC2 was designed and dramatically presented to the entire QSM community during the annual ISMRM meeting in 2019. The submissions were evaluated and results were presented at the 5^th^ International Workshop on MRI Phase Contrast QSM in Seoul.

In the remainder of this manuscript, we describe the challenge design rationale, provided data, evaluation criteria and results of the challenge.

## Methods (Challenge Design)

### Rationale

RC2 utilized a phantom with known ground-truth susceptibility, which was derived from in vivo data through MR physics simulations^22^. While addressing shortcomings of RC1, attention was paid to ensure that using phantom data did not cause new weaknesses. To avoid promoting piecewise smooth/contrast solutions, the susceptibility ground-truth included physiological texture and realistic variations within structures.

A realistic numerical phantom made it possible to synthesize gradient echo acquisitions, which provided local field map estimates without background field remnants. Inconsistencies between the simulated data and known ground-truths were restricted to complex Gaussian noise and intra-voxel dephasing effects, simulated by down-sampling from high-resolution 0.65 mm to lower-resolution 1 mm isotropic voxels^22^ via k-space cropping^7,23^. Due to the modular design of the simulations, other effects such as realistic background fields and shimming may be incorporated in a future challenge design. To evaluate the results, global and ROI-based metrics based on the root mean squared error (RMSE) were considered. The suitability of these metrics was corroborated by inspecting their correlations with other metrics (such as the Structural Similarity Index Metric (SSIM) and others) and a visual assessment.

The RC2 was designed to take place in two stages. Stage 1 assessed the reconstruction performance of the algorithms in a realistic setting where a ground-truth is not available. Participants had to determine optimal algorithmic parameters based on visual or numerical considerations (such as the L-curve) in the absence of a ground-truth. In Stage 2, given that ground-truth was available, participants could decide to numerically optimise their reconstruction algorithm with respect to one or all the metrics. The difference in performance between both stages was expected to provide insights into the ability to identify the optimimcal algorithmic parameters in the clinical settingground-truth.

### Provided Data

Both in the **first and second stages, two datasets** were provided related to two different susceptibility models (namely SIM1 and SIM2)^22^. The main difference between both datasets was the presence of an intra-hemispheric calcification (in SIM2), and different levels of susceptibility contrast between tissues, as seen in Figure 1.

**Figure 1.**
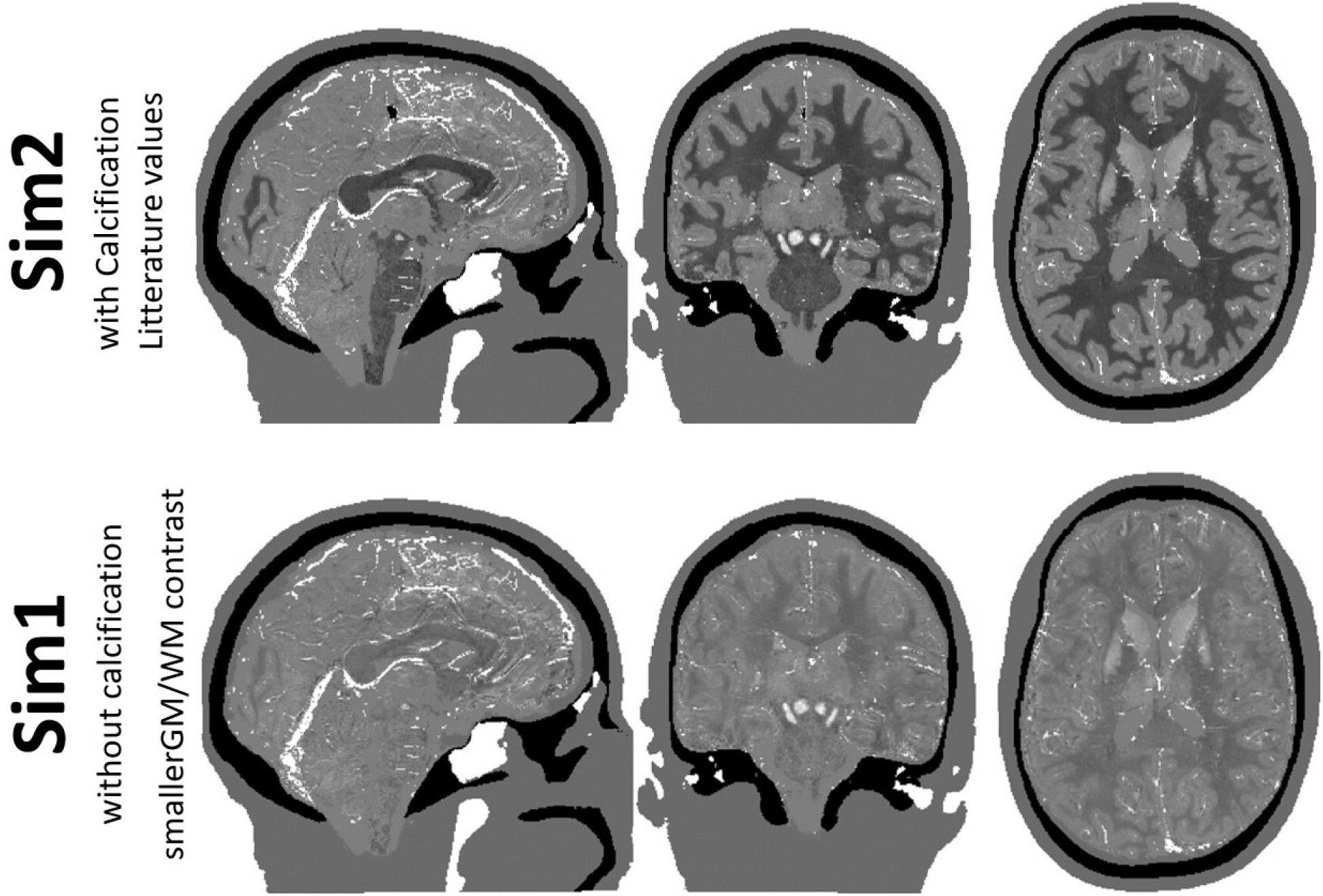
Ground-truth susceptibility maps used in both steps of the challenge. Sim2 presents a larger contrast between Grey and White Matter than Sim1, and includes a strong calcification. For RC2, all susceptibility values outside the brain mask were set to zero to remove background fields from the simulations.

As the focus of this challenge was the final dipole inversion step, only brain tissues were used to compute the field perturbations in the simulated MRI data. This ensured that there was no need for introducing a background field removal step.

Each dataset consisted of gradient echo magnitude and phase data simulated with the following parameters: TR=50ms; TE1/TE2/TE3/TE4=4/12/20/28ms; α=15°, FOV = 164×205×205 mm^3^ and 0.65mm^3^ isotropic voxels. Gaussian noise was added to the complex data at the same level for both datasets with peak SNR of 100 in Stage 1. For Stage 2, two datasets were generated, with peak SNRs of 100 and 1000, namely SNR1 and SNR2 respectively, for each susceptibility model (SIM1 and SIM2). The same k-space cropping approach^22^ was used to down-sample the high resolution (0.65mm to 1mm) complex signal, ground-truth susceptibility maps and segmentation labels. In the case of the susceptibility maps, the sharp edges between structures as well as the orders of magnitude larger susceptibility differences between air/bone and tissue resulted in severe Gibbs ringing artefacts, which were removed using sub-voxel shifts^22,24^. This methodology was repeated in all three spatial directions to ensure no Gibbs ringing remained.

Field maps were provided to be used at the discretion of the participants, by taking a magnitude and TE-based weighted average of three phase differences^23,25^ (TE_4_-TE_1_, TE_3_-TE_2_, TE_2_-TE_1_). Additionally, a mask corresponding to the brain region where the QSM reconstructions would be evaluated was provided. Figure 2 shows an overview of the MRI data provided at the two stages.

**Figure 2.**
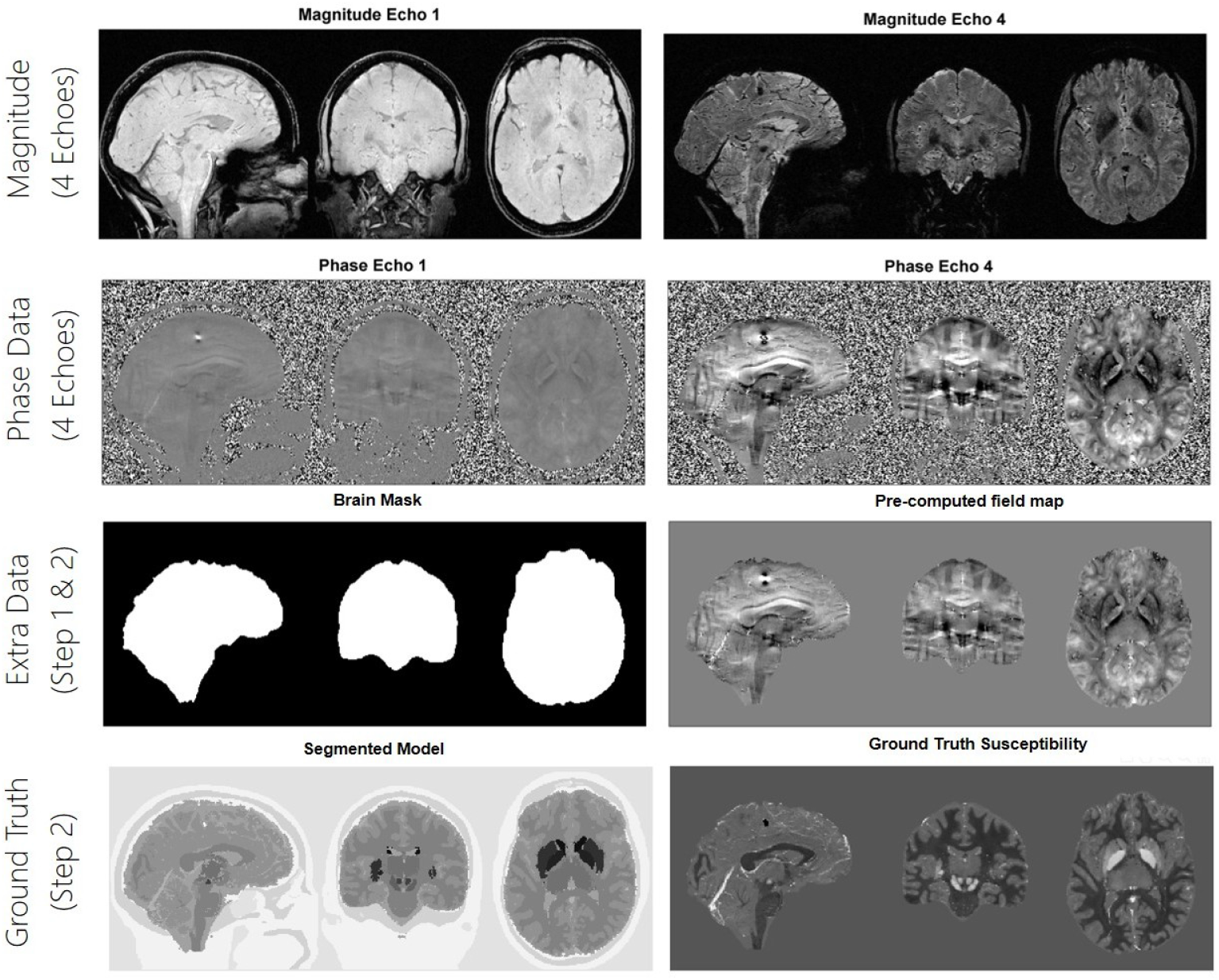
Data provided to participants in Step 1 and 2. In the first step, participants were provided with 1mm isotropic whole brain multi - echo magnitude and phase data (1st and 2nd row), simulated in the absence of background fields. A precomputed brain field map was provided as well as a mask reflecting the region where the scripts would be evaluated (third row). In step 2 the ground-truth maps were given as well as the segmented model of the brain (4th row) that was needed to compute the various specialised metrics.

In Stage 1, participants were asked to use the same processing pipelines for the two datasets (including regularization parameters or regularization optimization). Iterative and closed-form algorithms were able to optimise once again their algorithm parameters for Stage 2. DL algorithms were able only to modify certain parameters such as epochs, batch size, etc., but they were not allowed to modify the architecture of the network or to incorporate the groundtruths into the training set.

### Announcement and Participation

Data and instructions for participation were disseminated on a publicly accessible website (http://qsm.rocks) following the 2019 Annual Meeting of the ISMRM in Montreal^26^. The deadlines for submission of solutions for participation in Stages 1 and 2 were originally set as June 27th and August 27th, 2019, respectively, and extended to August 2nd and September 1st, 2019, respectively.

### Data Management and Evaluation

The challenge evaluation was designed to allow a fully blinded analysis of submitted solutions. While participants were required to submit personal as well as detailed algorithmic information along with their susceptibility maps at the time of participation, we ensured that personal and algorithmic information was not accessible to the analysis team. Until the announcement of the challenge outcome, only one committee member (F.S.) had access to the identifying information of the participants. Participants were asked to provide a long-form algorithm name as well as an acronym, and a self-chosen random (blinded) 10-characters identifier for the submitted solution. Please see the Supplementary Information for a detailed description of the required form fields.

Participants were asked to include their solution in a compressed zip-archive that was named after the self-chosen identifier and upload it through an interface to a file server (Nextcloud; hosted by F.S.’s institution). No further identifying information was supposed to be contained in the archive. Until the announcement of the challenge outcome, only one committee member (F.S.) had access to the identifying information of the participants. Read-only access to the file server containing all submissions (but not the online form data) was given to the analysis team after August 2^nd^ for Stage 1 data and September 1^st^ for Stage 2 data. Algorithmic information was only shared with the analysis team after all metrics had been computed and the winners for each category were already established, on September 16^th^ 2019.

### Metrics

If the input phase data contains only information compatible with the magnetic dipole convolutional model, it is to be expected that most of the global metrics tend to produce the same optimal reconstruction parameters for a given algorithm^16^. Phase data inconsistencies or external contributions lead to a disagreement of the optimal parameters^16^, as shown in RC1. Given that RC2 consists of phantom-based forward simulations, to avoid unnecessary complexity in the challenge design (winning categories), only RMSE-based metrics were chosen to officialy evaluate the global performance of the submissions.

In addition to global error metrics, we included three ROI dependent error measurements (Tissue, Blood and Deep Grey Matter), with the aim to provide an assessment more closely related to clinical needs, such as those found in QSM-based oximetry^27,28^ or the study of deep brain structures.

The metrics chosen for evaluation were:

- NRMSE: Normalized RMSE, inside the ROI. Normalization is performed by the L2-norm of the respective ground-truth.
- dNRMSE: Some algorithms are known to produce underestimated results. To address this issue, we included a data demeaned and detrended RMSE score.
- dNRMSE Tissue: dNRMSE specific to White Matter and Grey Matter tissues.
- dNRMSE DeepGM: dNRMSE specific to Deep Grey Matter structures.
- dNRMSE Blood: dNRMSE for blood regions (effectively it was a dilated version of the vein mask).
- Deviation From Linear Slope: Absolute error of the slope, derived from the demeaning and detrending process.
- Calcification Streak: Error metric based on the standard deviation inside a square neighborhood surrounding the calcification, in the difference map.
- Deviation From Calcification Moment (CalcificationError): Error in the quantification of the total moment of the calcification, defined as the volume of the reconstructed calcification multiplied by its mean susceptibility.

Further details are described in the Supplementary Information. To provide additional validation of the measured metrics and the winning categories of the Challenge, we also calculated SSIM, (both with the standard formulation, and a QSM-specific formulation called XSIM^29^), the High Frequency Error Norm (HFEN)^11^, the Correlation Coefficient (CC), Mutual Information (MI), Mean Absolute Difference (MAD), and the root mean squared error of the gradient (first derivatives) domain (GXE)^16^.

Scores were averaged across SIM1 and SIM2 submissions to reduce the complexity of the analysis.

### Visual Rating

A visual rating scheme was designed to complement the numerical assessment of the submitted solutions. The rating was performed by each of the challenge committee members (authors) individually. A MATLAB graphical user interface was created for this purpose (Supplementary Information Figure S1).

The submissions from Stage 1 of the Challenge were presented individually and in random order, different for each rater. After rating for one category, the next category was rated, using the same random order. Images for Sim1 and Sim2 were rated together, and the worse of the two submissions was used to determine the score. Each submission was shown for three fixed orthogonal slices. This set of slices covered: large veins, major deep gray matter region and the calcification region. This restriction was done for three main reasons: (i) minimise the transferred data across raters and memory requirements; (ii) speed up the visual rating for the raters, allowing to quickly navigate through various submissions and previously given ratings to ensure consistency; (iii) ensure that the evaluation was based on the same aspects of the image reconstruction increasing the consistency across ratings.

The ground-truth was not shown alongside the submitted solutions, but shown as if it were a submission itself. For the final score, the ratings of all raters were averaged.

Scores from 0 (best) to 3 (worst) were given depending on the artifact level in three distinct categories (Streaking, Unnaturalness, and Noise), which are described in the Supplementary Information. It is to note that Visual assessment is not a comparison between QSM reconstructions and the ground-truths, but a quality assessment of the naturality, and lack of noise or artifacts. A high scoring image (by visual evaluation) may contain significant errors such as “hallucinated” structures or misplaced sources. This is of special concern in the case of evaluating DL algorithms. To account for this, in addition to these three categories, a binary rating was performed based on the difference map obtained by subtracting the ground-truth from the submission. These are referred as visual Discrepancy (White Matter/Gray Matter, Deep Gray Matter, and Veins) metrics.

### Data Preprocessing and Quality Checks

Prior to the evaluation, a check on the dimensions of the reconstruction and global scaling was performed. When clear mismatches existed, authors of the submissions were invited to resubmit their solution. The resubmissions were manually checked to ensure that only rescaling or spatial shifts had been applied to the new solution.

Stage 1 submissions were awarded in two categories: 1) NRMSE performance, and 2) “Robustness”, which counted how many appearances an algorithm had in the top 5 of any metric. Honorable mentions were awarded to the best performing algorithms in Stage 2.

## Results

### Participation and Submission Statistics

We received 98 unique submissions for Stage 1. Of those submissions, 47 were submitted to Stage 2 as well (excluding resubmissions). An extended description of the number of downloads and participating countries is provided in the Supplementary Information.

The majority (85%) of the submissions used an algorithm that was described either in a published journal paper or conference abstract^14,15,17–21,30–56^ (Supplementary Information Table S1). 15% of the submissions were not yet published. Further details regarding submitted algorithms are presented in the Supplementary Information.

### Open science

Source code of the QSM algorithms was publicly available for 20% of the submissions (Supplementary Information Table S1). Participants agreed to make the algorithm code available after the challenge for 43% of the submissions and they stated they would “maybe” make the code available for 37% of the submissions. None of the participants refused to make the code available on request.

### Results of Stage 1

Figure 3 shows how the NRMSE differed between both simulations, as a function of the average NRMSE, for different algorithm types. Overall, algorithms showed lower errors for SIM2 with the exception of most DL-based methods, which had a similar performance in both simulations. It is important to note that SIM2 has a higher contrast between grey and white matter as well as a diamagnetic calcification, which leads to a higher normalization factor in NRMSE calculation (SIM2:31.4 vs SIM1:17.4), thus lower NRMSE scores were expected for SIM2. Interestingly the best performing deep learning method (FINE)^21^ had a notorious different performance on both simulations.

**Figure 3.**
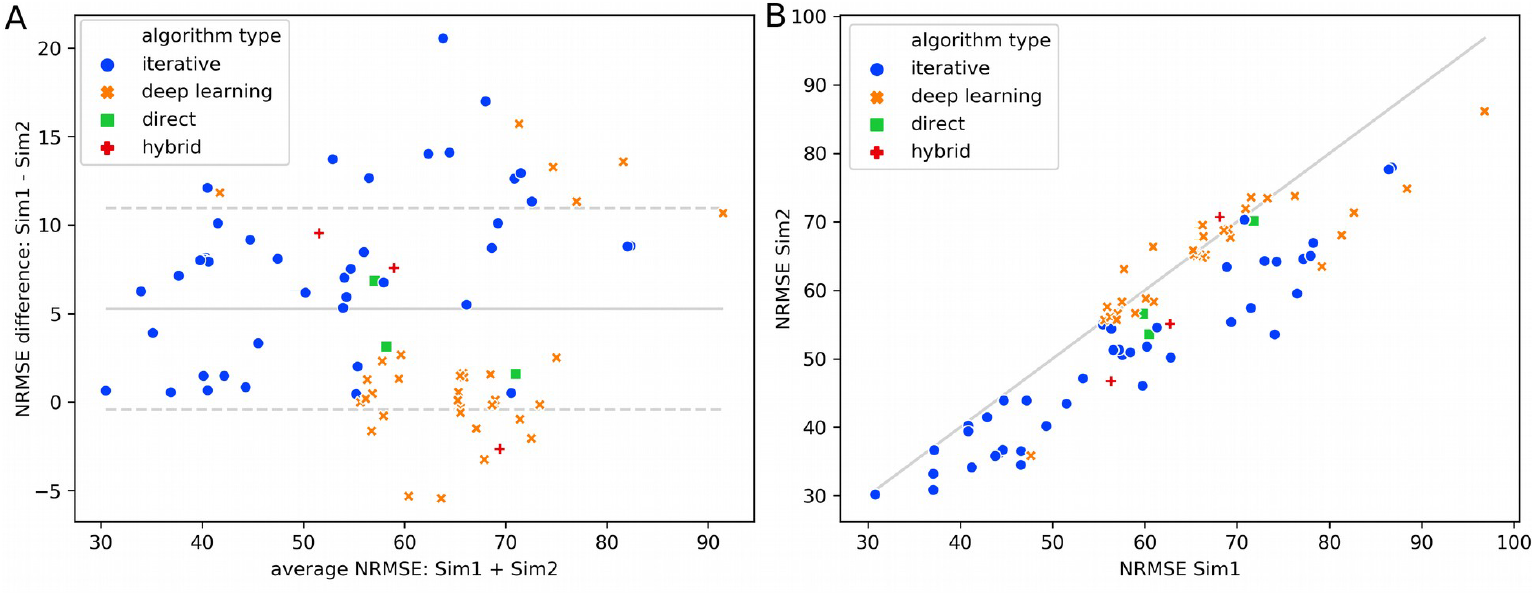
Analysis of Sim1 and Sim2 submissions: A) Bland-Altman plot, and B) Sim1 vs Sim2 NRMSE scores. Color codes show different algorithm types. Deep Learning approaches performed worse on average (higher average NRMSE), but more consistent across Sim1 and Sim2 (lower difference in NRMSE) compared to spatial-domain iterative approaches. In general, no systematic differences between Sim1 and Sim2 was noted, and therefore further analysis was performed on averages across Sim1 and Sim2.

Figure 4 shows a correlation matrix for the various metrics, both derived visually and numerically. Only submissions with NRMSE<80 were included (83 submissions). RMSE-based metrics were highly correlated (R≥0.96). RMSE-based metrics were also highly correlated with additional global metrics such as SSIM (XSIM variant), HFEN, CCMI and MAD (Supplementary Information Figures S2-S3). Correlations between analytical metrics were significantly higher than between visual metrics. Visual metrics also correlated fairly well with RMSE metrics (r=0.66 for the mean visual score). Note that the (Un)naturalness metric had the highest correlation with the global, deep grey matter and tissue RMSE metrics. This is particularly relevant because in a normal scenario, in the absence of a ground-truth, (un)naturalness may be the criteria to choose one particular reconstruction pipeline. The visual streaking rating had its highest correlation with the Calcification streaking metric. Visual discrepancy correlated fairly well with calcification metrics (r>0.45) and RMSE metrics (r>0.40). Veins discrepancy correlated with r=0.44 with the Blood RMSE. Visual Streaking correlated very well with the Calcification dissimilarity (r=0.81). Visual noise was not significantly correlated with most error metrics (except GXE, which correlated poorly with other metrics). This is to be expected as in the definition of the visual noise metric, over regularized solutions (that tend to have higher RMSE) could achieve high ratings (note that the ground-truth was not top-ranked in this metric). The ground-truth map ranked 1st (together with 6 other submissions) for “Visual streaking”, 1st (together with 2 other) for “Unnaturalness” (most natural) and 7th (together with 15 more) for the (lowest) visual noise level. For the visual metrics, inter-rater correlations varied from 0.30 to 0.80 for the “Unnaturalness” metric (the most subjective metric) while for the streaking and noise metrics they varied from 0.46 to 0.83. The Dissimilarity metrics had broader inter-rater variability (0.06 to 0.81) because of their binary nature.

**Figure 4.**
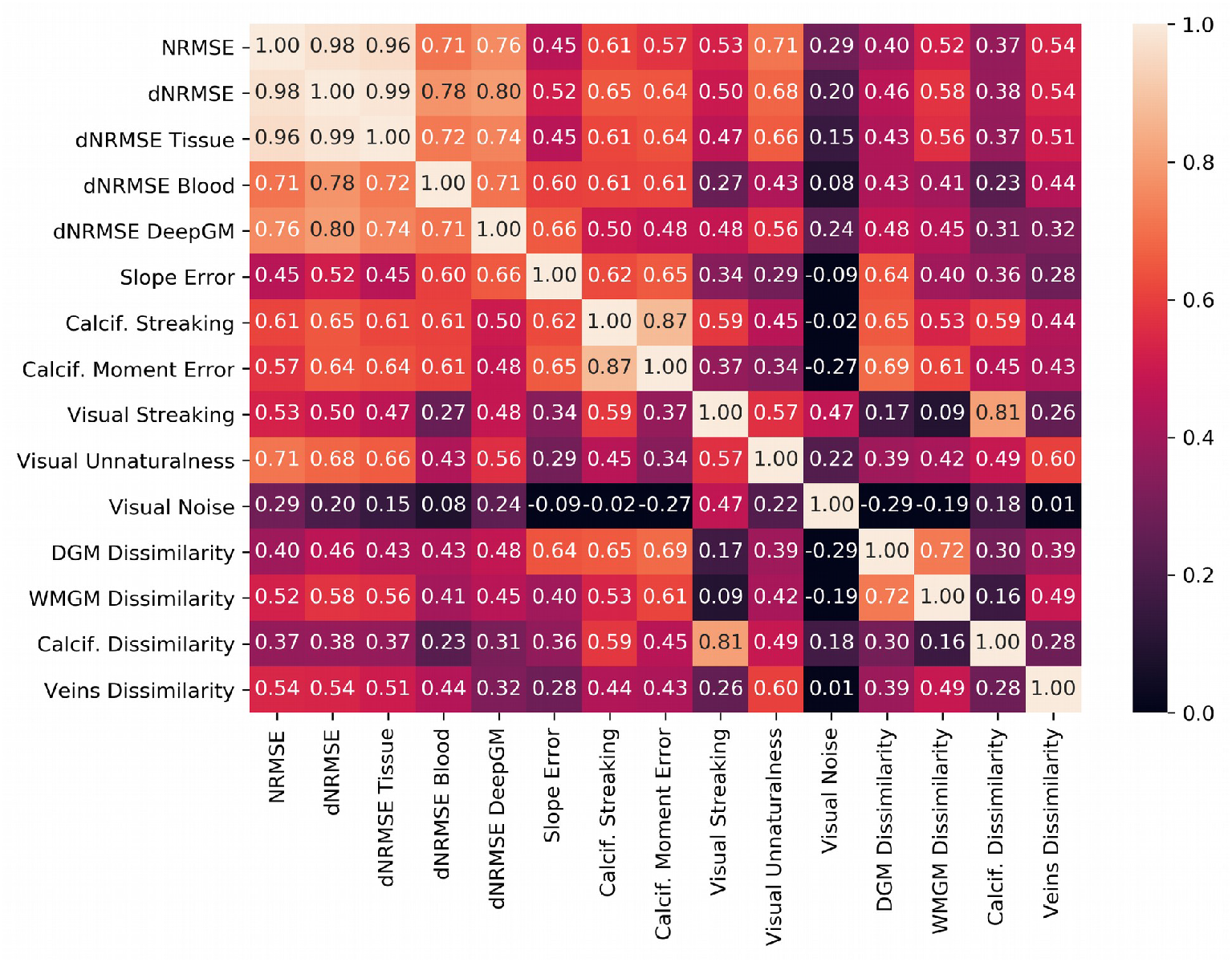
Correlation between metrics, for all Stage 1 submissions with RMSE < 80.

Plots with the scores of the top 20 scoring submissions for each algorithm type are presented in Figure 5. Overall, iterative algorithms performed better than direct and DL algorithms, in all metrics. The performance of DL algorithms was more similar to iterative algorithms in visual analysis, and was considerably worse in the NRMSE. A similar analysis is shown in Supplementary Information Figures S4-S6, where the difference in performance is shown depending on whether the supplied frequency map was used as input or the full multi-echo dataset, and similarly for the case of using magnitude information. While using the magnitude information provided a small advantage, submissions using the multi-echo data (either to estimate a new field map, or as part of the algorithm’s input/functional) showed on average a clearly better performance across all metrics. Most submissions that did not use the magnitude information used the provided frequency map. Scatter plots for additional metrics are shown in Supplementary Information Figures S7-S8.

**Figure 5.**
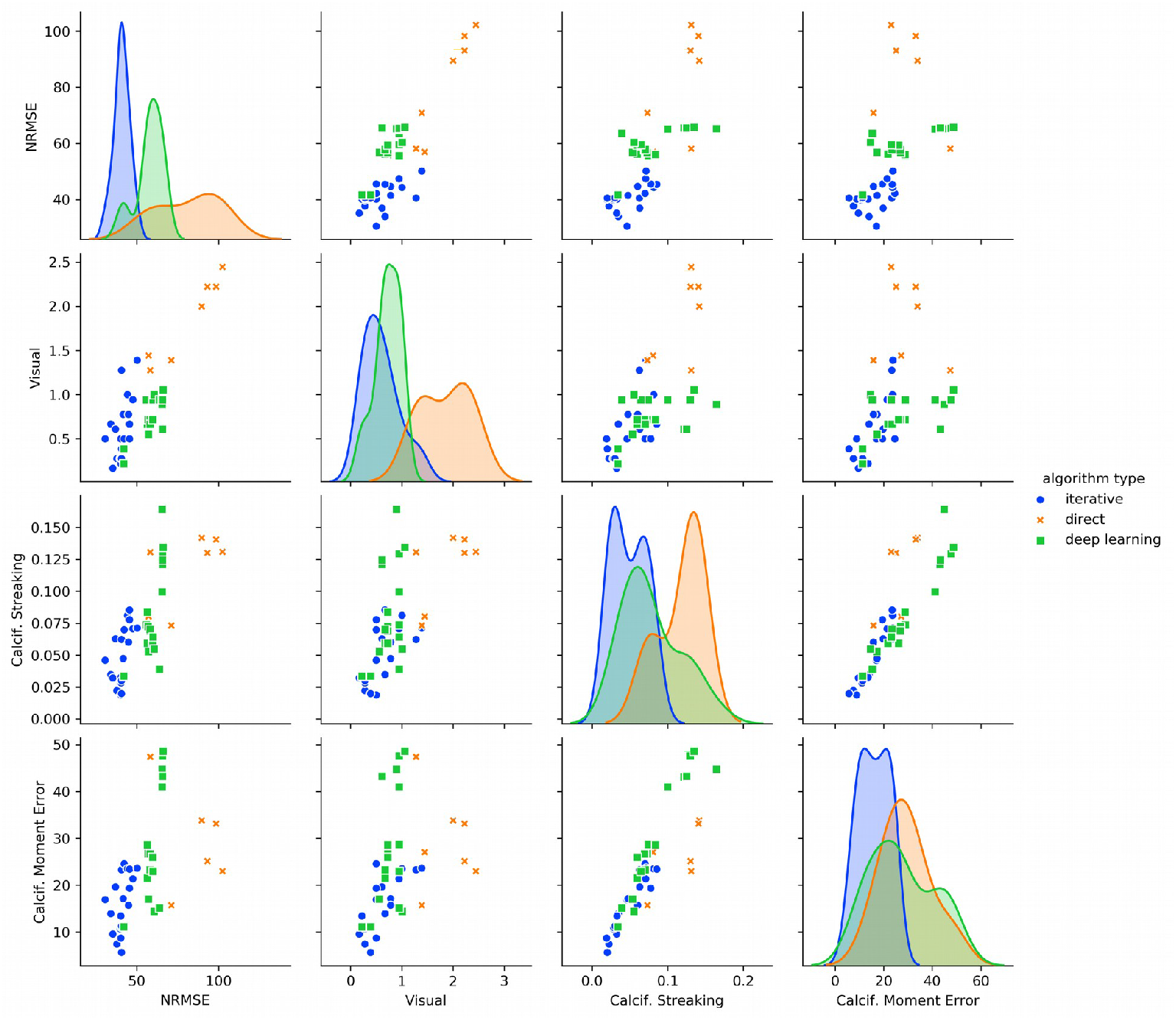
Scatterplots between selected pairs of metrics showing the top 20 NRMSE (Stage 1) submissions in each algorithm class (shown as different colors, see legend). The diagonal shows estimated histograms for each metric. In general, the top 20 solutions using iterative methods show lower error metrics than deep learning based methods, which in turn show lower error metrics than direct inversion methods.

Top 5 scoring results in the NRMSE metric and the Honorable Mentions (discussed in the next section) are shown in Figure 6. Coincidentally, the top 5 NRMSE were also the top scoring algorithms in the “Robustness” category. The winner of the Best NRMSE category was submission 5e0YBJKZv1 (wL1L2 algorithm^48^, from Pontificia Universidad Catolica de Chile), which used a combination of L1-norm^49^ and L2-norm data fidelity terms and an R2* weighted TV regularization (FANSI toolbox^36^). The winner of the Robustness category was submission WgpBiTiZw9 (mcTFI algorithm^50^, from Cornell University), which used an extended functional that included all individual echoes, a preconditioner derived from R2* data, and a morphologically enforced TV regularization (MEDI^33^ toolbox). Metric scores for the top 10 submissions in these two categories are shown in Table 1. The performance of the top-scoring algorithms using the NRMSE, CalcStreaking and CalcificationError metrics is visualized comparatively in Figure 7.

**Figure 6.**
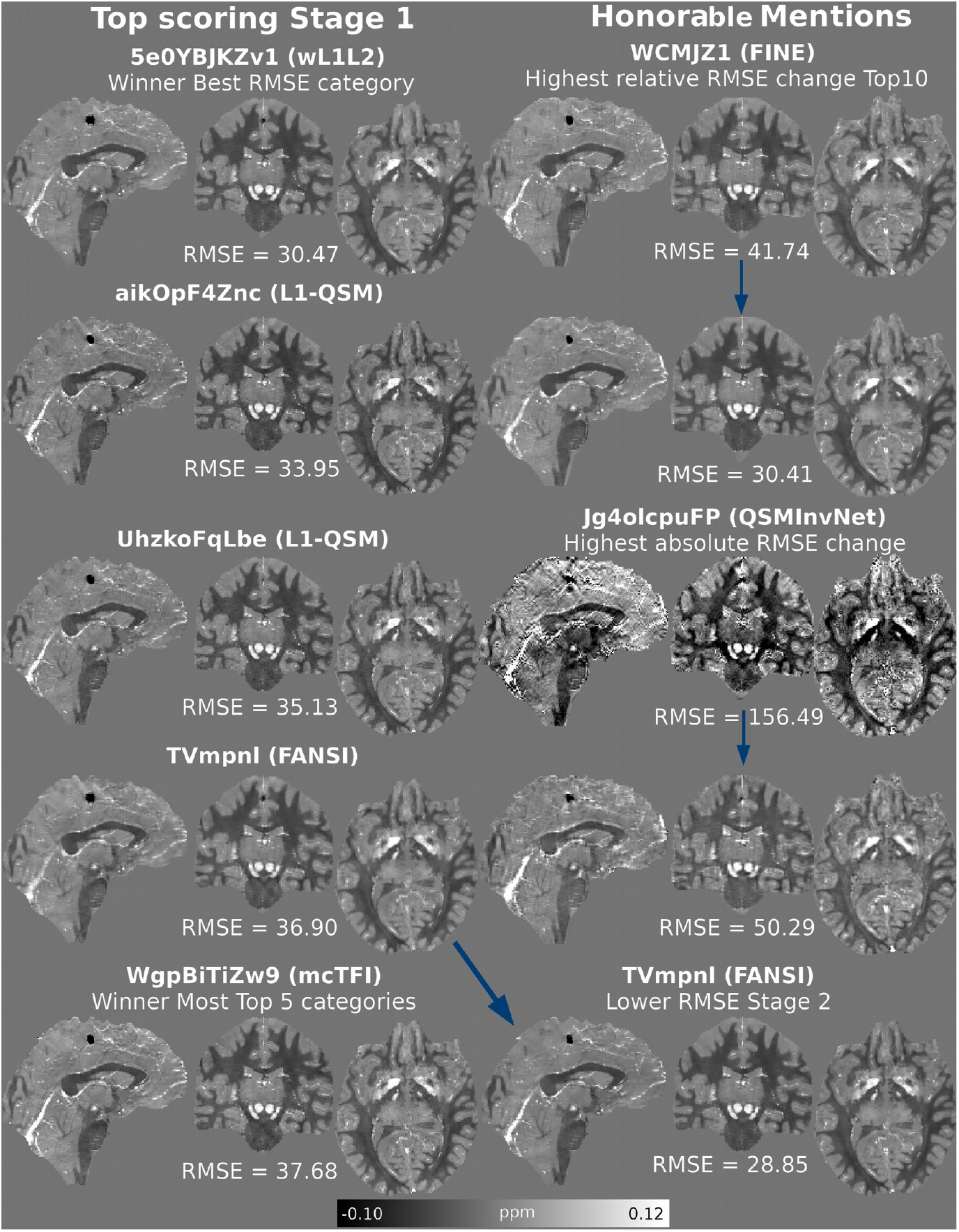
Left: Top 5 RMSE scoring submissions, in descending order (top to bottom, with wL1L2 the RMSE winner). They were also the top scoring submissions with most appearances in the top 5 of all metrics (robustness category). The mcTFI algorithm was the robustness winner, with 7 points. All other shown algorithms tied with 4 points. Right: honorary mentions, based on the RMSE performance in Stage 2.

**Table 1.**
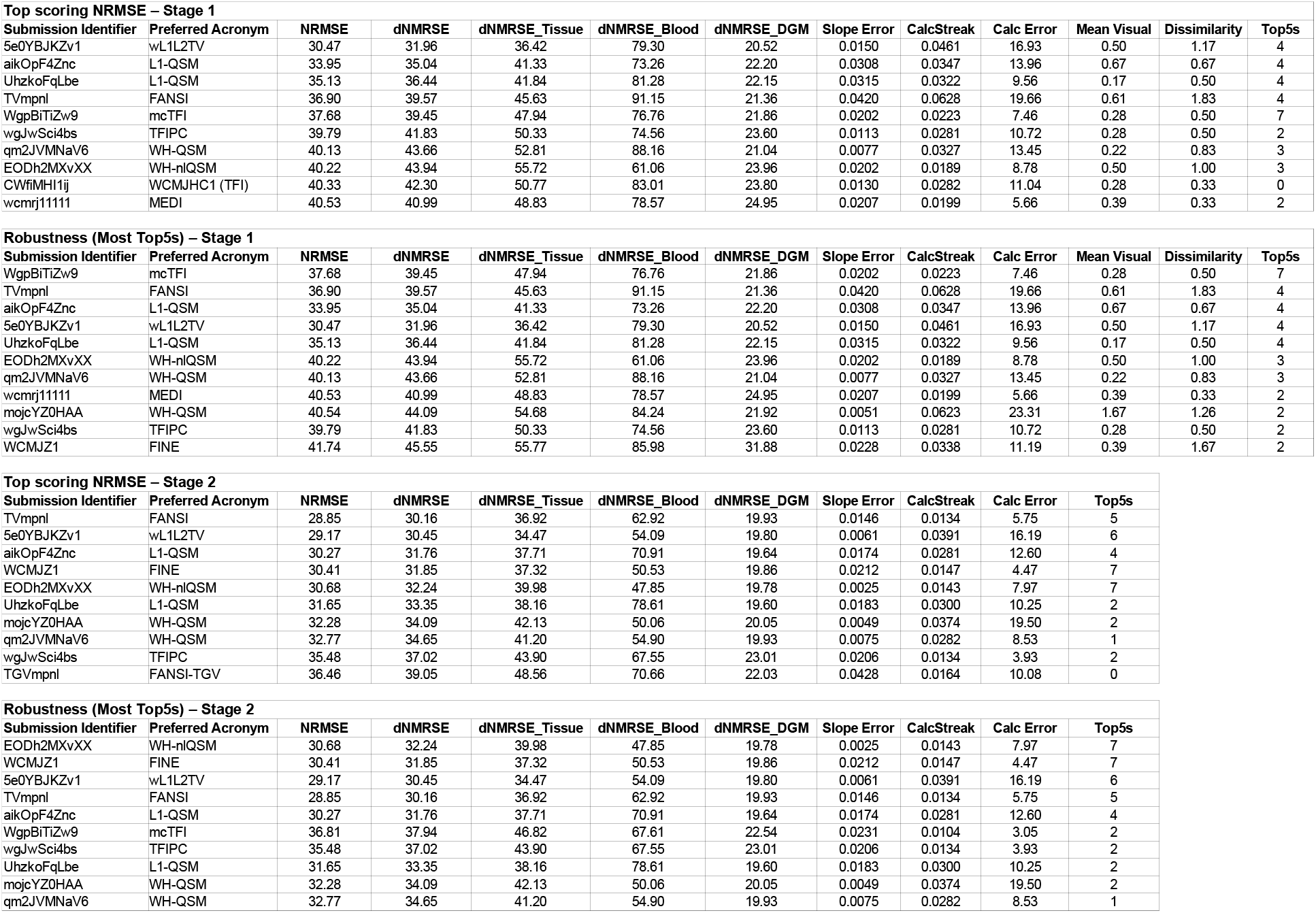
Metric scores for top 10 scoring results in the awarded categories (NRMSE and number of appearances in the top 5 scoring results for each measured metric, or “robustness”). Top half shows the results for Stage 1 submissions, and bottom half for Stage 2 submissions.

**Figure 7.**
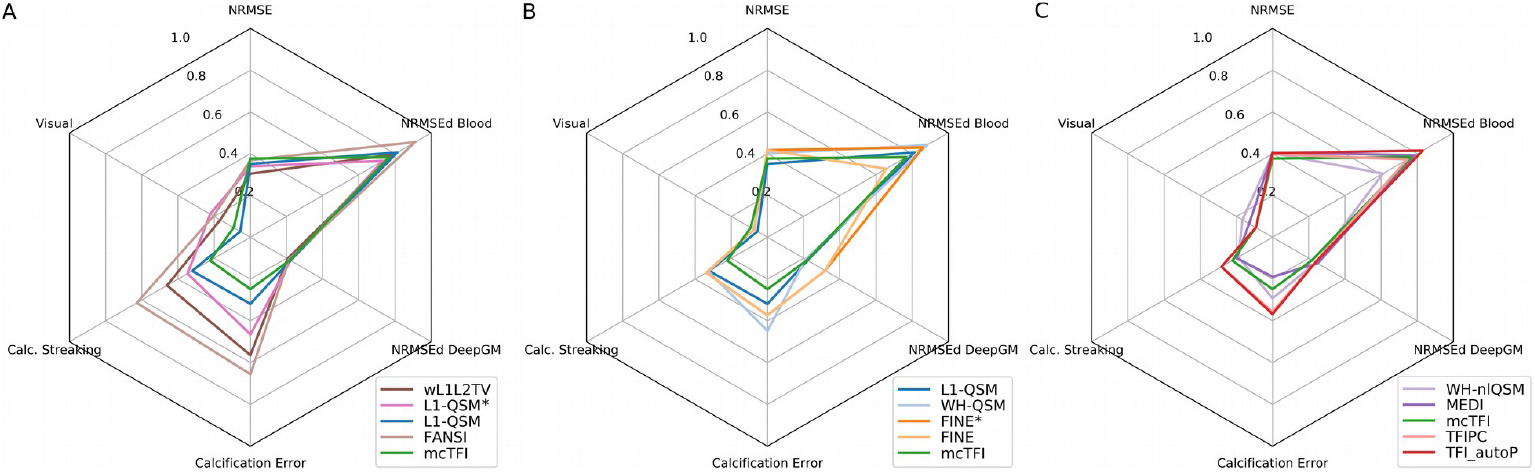
Spider plots showing metric scores for top scoring submissions. (A) Top5 submissions sorted by NRMSE. (B) Top5 submissions by Visual Rating (average). (C) Top5 submissions sorted by Calcification Streaking. Scores where normalized by the following factors, for displaying purposes: RMSE-based metrics: 100.0, (mean) Visual: 3.0, Calc. Streaking: 0.1, Calcification Error: 30.0. Duplicated algorithm acronyms correspond to multiple submissions with different inputs or parameters.

### Results of Stage 2

With a few exceptions, rankings for Stage 2 remained similar to those in Stage 1. As shown in Figure 8, a few DL submissions performed worse in RMSE metrics, with slight improvements depicting the calcification. Calcification-related metrics showed little or no improvement for most submissions, with some algorithms performing considerably worse. Similarly, some DL submissions had inferior RMSE performance with improved SNR, whereas most algorithms performed better (Figure 9). The calcification-related metrics seemed to be less dependent on the SNR, with mixed results across all algorithm types. Correlation and scatter plots for Stage 2 submissions are shown in Supplementary Information Figures S9-S12.

**Figure 8.**
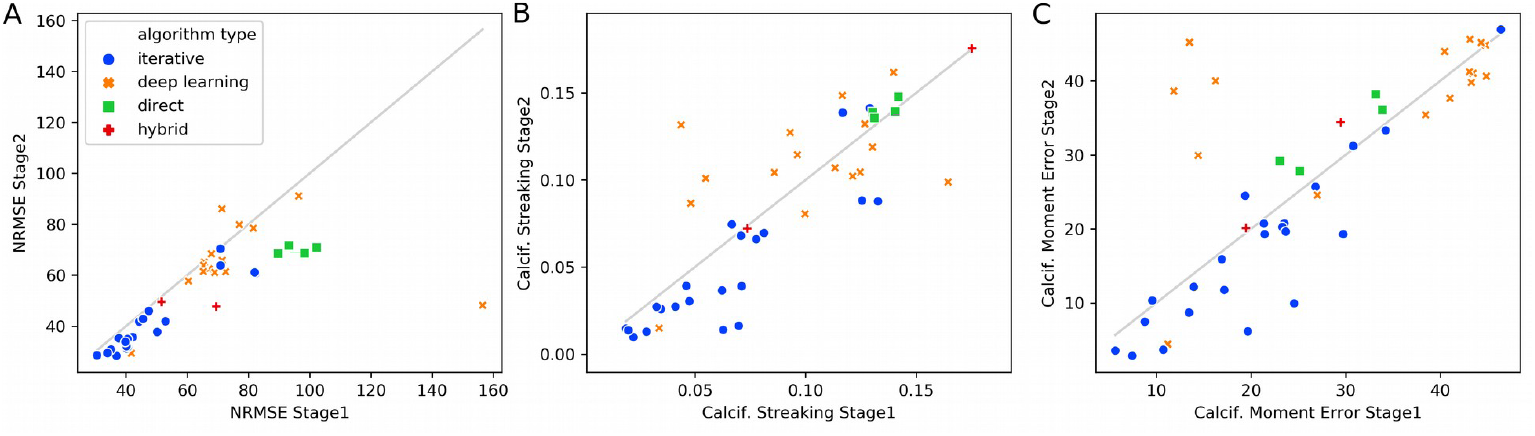
RMSE, Calcification Streaking, and Calcification Moment Error change between Stage 1 (horizontal axis) and Stage 2 (vertical axis). The grey lines indicate the “no changes” regime. Solutions over this line worsened their scores, while solutions below this line improved their results.

**Figure 9.**
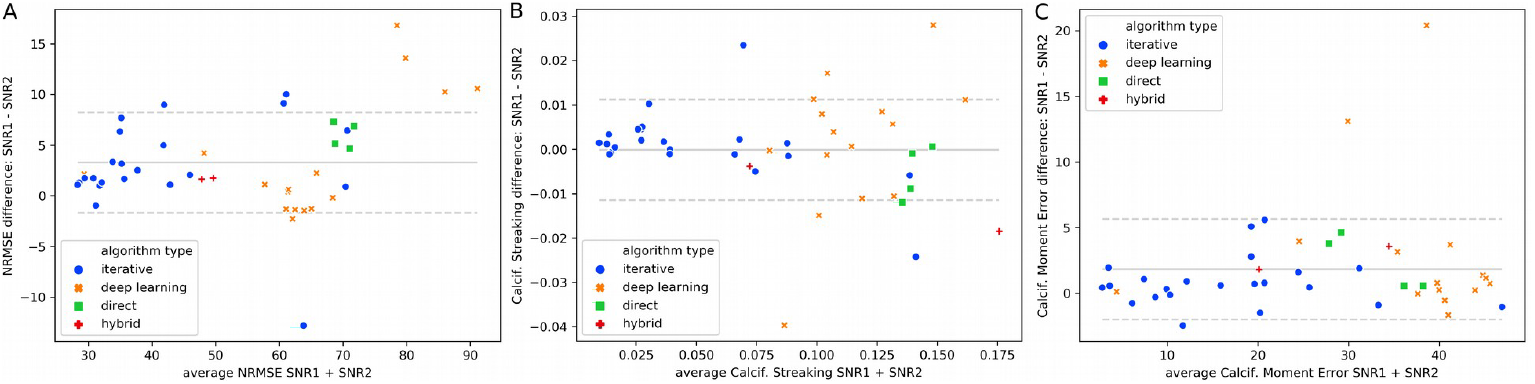
Bland-Altman plots comparing SNR1 and SNR2 Stage 2 submissions for the: A) RMSE, B) Calcification Streaking, and C) Calcification Moment Error metrics.

The submission with the best RMSE score (Honorable Mention) was TVmpnl (FANSI^36^ algorithm, from Pontificia Universidad Catolica de Chile), which used a nonlinear data fidelity term and TV regularization. The submission with the best relative NMRSE improvement (for top scoring algorithms) was WCMJZ1 (FINE^21^ algorithm, from University of Cornell), a DL algorithm with an imposed fidelity term that modifies the pre-trained weights. This was also the overall best performing DL algorithm. Finally, the highest absolute RMSE improvement was performed by submission Jg4olcpuFP (QSMInvNet^30^ algorithm, from the Medical College of Wisconsin), which used a Nonlocal Encoder-Decoder convolutional network. Top scoring Stage 2 results for the RMSE and “robustness” categories are also shown in Table 1.

## Discussion

Compared with the 1^st^ QSM reconstruction challenge, the present RC2 was based on an entirely different concept. The availability of a ground-truth and its 2-stage-design allowed more in-depth insights and the usage of additional metrical analysis than with the in vivo dataset from RC1.

The analysis of the two datasets with significantly different SNR provided in the stage 2 showed that SNR had little impact on the ordering of the ranking (note the different scaling in fig. 9), suggesting that most algorithms behave in a similar way to different noise levels. Also, the usage of the two susceptibility models, with and without calcification, for which the participants had to submit the reconstructions with the same reconstruction parameters (although it was acceptable to submit multiple times with different parameters) did not have the expected impact. Most methods were able to deal with both datasets with similar performance (see Fig. 3).

Overall, taking NRMSE as the main metric for evaluation seemed justified, as this metric was highly correlated with all other global metrics. This might be a natural consequence of avoiding the presence of phase incompatibilities, which generate artifacts in the reconstructions. In the presence of such errors, different metrics promote different features or image properties, thus resulting in results with different algorithm parameters when they are used to optimize the fidelity/similarity with a ground-truth^16^.

For all metrics, iterative methods performed better in both stages. Metric scores for SIM 2 tended to be better than for SIM 1, mainly because of the increased WM-GM contrast. In particular, methods based on Total Variation^15,19,33,36,48-50^ (and derivatives such as TGV^35,36^) were the top-scoring algorithms. Direct inversion methods (TKD^55^ and Tikhonov^51,52^) had inferior performance. Interestingly and contrary to common expectations, QSM methods based on DL techniques were outperformed by “conventional” iterative methods. DL methods showed a large variance but overall performed significantly worse than iterative methods. An exception was a DL algorithm that used a physical model as the fidelity term, with similar performance to the top-scoring iterative method group^21^. Noteworthy, some DL algorithms performed worse in Stage 2. The underlying reasons for sub-par performance of DL methods could be that the networks failed to generalize well to the RC2 dataset since this fell out of the distribution of susceptibility maps they were trained on, and that the training data might be coming from a model with large data inconsistency such as COSMOS^13,17^. Poor generalization could also be the reason why some DL methods yielded worse results when using higher SNR data, and why they performed considerably worse in the calculation of the calcification moment. Also, the non-linear nature of noise^57^ might have been neglected in the DL methods either in the loss function during training, or in the forward model during inference^58^. Furthermore, DL methods are quite novel compared to the established iterative methods with more than 10 years of development and refinement. However, besides the aforementioned factors also other aspects beyond our current intuitive understanding might have contributed and require systematic analysis.

Overall, errors in the estimation of susceptibility values in the vessels were notoriously high, although they correlated very well with other global metrics. It is unclear if this is a cause of intra-voxel dephasing, in such small-scale and high-contrast regions, or an intrinsic problem in QSM methods to estimate large dynamic range data. While previously reported in the literature, most algorithms showed no significant understimation of susceptibility values in this challenge. Demeaning and detrending did not change the results in a significant way. Whether the issue of underestimation has been resolved in current QSM algorithms or underestimation is a consequence of phase inconsistencies not modeled in the RC2 remains to be investigated.

The 5 top-ranked algorithms in the category RMSE also demonstrated similar error metric results in Blood and DeepGM dNRMSE (Figure 7 and Table 1), which is expected given the high inter-metric correlations (Figure 4). Especially the error in DeepGM showed a low variance for those algorithms, which provides confidence that the susceptibilities of those iron-rich nuclei can be consistently compared between different algorithms and studies.

Generally, the submitted results allow the conclusion that multi-echo fitting or using all the echoes in the data fidelity term produced better results than using the provided field map, which is calculated from a phase difference method. Using the whole echo train improves the SNR and reduces the presence of artifacts in the vessels and the calcification. It is not clear if this conclusion can be generalizable to the case where a significant initial phase shift exists at TE=0. Using an extended data fidelity term that incorporates all the echoes in the forward model (as in mcTFI^50^) also provided better solutions to the problem of intra-voxel dephasing in voxels surrounding the calcification.

The calcification introduced in SIM 2 basically split the algorithms in 2 domains, those yielding the typical streaking artefacts and another group of algorithms handling such discontinuities better by the utilization of preconditioners or voxel rejection strategies. A related question is whether models including estimations of the background field performed worse in this case. As expected, since no background fields were simulated, additional and unnecessary background field removal yields worse results. One example is the case of FANSI^36^ vs WH-QSM^15^, where the latter includes background field remnants in its data model. The calc-streaking metric presented an evaluation of the standard deviation in the surroundings of the calcification. It tended to promote solutions with less streaking, but it was not robust. Calculating the error in the calcification moment seemed to be a more robust metric, but it has a lower correlation with the visual assessment (r=0.22 for the mean visual score).

Using the magnitude information provides advantages, although it is not clear whether this is more relevant as a SNR-based data-fidelity weighting, or as a local constraint in the regularization (i.e. morphological consistency between susceptibility maps and the magnitude).

Rather a conceptual decision than a limitation was the utilization of synthetic GRE datasets, which were derived from several high resolution 7T scans^22^. Although the generation of these datasets incorporates a variety of processing steps and emulates conditions close to real-world 3T brain scans, it is acknowledged that in vivo GRE acquisitions might yield different results, but would also preclude the availability of a ground-truth susceptibility. Although the provided ground-truth scored as a natural image in the visual rating when compared to the remaining reconstructions, an experienced viewer might notice that it lacks texture in regions such as the thalamus, ultimately still appearing as a piecewise smooth model. This could have had an impact in the observation that all the top-rated RMSE solutions were TV-based. Researchers wanting to further develop their methods are encouraged to explore new realizations of the digital phantom and simulation data using the toolbox provided in the part 1 of this manuscript which could have a more natural appearance. Another issue is the inclusion of susceptibility anisotropic and microestructural effecs, along with phase inconsistencies arising from flow artifacts and other effects. Further studies or challenges should be able to assess the robustness of inversion algorithm to this type of effects, more closelly resembling in vivo clinical settings and not an ideal scenario such as the one presented here.

## Conclusions

RC2 constituted a 2-stage challenge design based on synthetically generated brain GRE data, and yielded novel insights, which may not be obtained using an in vivo GRE acquisition. It aimed to overcome the shortcomings of the previous challenge, such as background field remnants, low SNR and the absence of a reliable ground-truth. Utilizing the RMSE as a fidelity metric in RC2 was successful indicated by a high correlation with other global metrics and the visual assessment. Iterative methods had generally better performance than DL and direct inversion methods. Incorporating the information from all the echoes and magnitude images yielded better metrics. While the synthetic phantom allowed for evaluating the performance of the algorithms in a challenging scenario with calcification, its design remains modular enough to incorporate additional considerations such as anisotropy and background fields. The data and exemplar code are publicly available, which will facilitate the development and benchmarking of future dipole inversion algorithms.

## Data Availability Statement

The code and data of this QSM reconstruction challenge are openly available in: https://surfdrive.surf.nl/files/index.php/s/uTvrxJ5NELbnGa5

Original anonymized cvs files used for Workshop analysis and original submission forms: https://doi.org/10.5281/zenodo.3687196

All original Stage 1 submissions: https://doi.org/10.5281/zenodo.3687341

All original Stage 2 submissions: https://doi.org/10.5281/zenodo.3688702

Figure creation code:

https://doi.org/10.5281/zenodo.4117549

The phantom/acquisition generation toolbox^22^:

https://data.donders.ru.nl/login/reviewer-113366422/QRFk431i299BX-8bcY6ta6nQPP-MqSzG0DTIhkJrqBs

## Acknowledgements

We are grateful for extensive discussions at the Electro-Magnetic Tissue Properties ISMRM study group meeting. We also thank Dr. Karin Shmueli, Dr. Pinar Ozbay and Dr. Cristian Tejos for their valuable input in early design stages. CL was supported by the Austrian Science Fund (FWF grant numbers: KLI523, P30134) and BioTechMed-Graz, JPM received support from the Nederlandse Organisatie voor Wetenschappelijk Onderzoek (NWO - Grant/Award Number: FOM-N-31/16PR1056). CM received support from FONDECYT 1191710, PIA-ACT192064 and the Millennium Science Initiative Program – NCN17_129, of the National Agency for Research and Development, ANID. BB was supported by the NIH (R01 EB028797, U01 EB025162, P41 EB030006, R01 MH116173, U01 EB026996) and Siemens Healthineers. FS was supported by the National Center for Advancing Translational Sciences of the National Institutes of Health under Award Number UL1TR001412. The content is solely the responsibility of the authors and does not necessarily represent the official views of the NIH.

## Supporting Information

### Data Management supporting details

To download the data for Stage 1, participants had to enter their email address. This step enrolled them into the RC2 email distribution list through which all challenge-related communication with the participants was conducted.

### Submission of solutions for Stage 1

Submission of solutions for Stage 1 required the submission of an online form (Google Forms) that collected identifying information of the participant as well as detailed information about the applied QSM technique. Personal information included email address, full name, institution, department, lab name and head, and address. Algorithmic information included the following mandatory questions:

- Algorithm-type (one of: closed-form direct solution; inverse filtering (TKD-like); spatial-domain iterative reconstruction; deep learning; hybrid (multiple categories); or a free-text field)
- Does your algorithm incorporate information derived from magnitude images? (yes/no)
- Regularization terms (None or does not apply; L0; L1; L2; Wavelets; total-variation; generalized total variation; and a free-text field)
- Did your algorithm use the provided frequency map or the four individual echo phase images? (frequency map; individual echo phase images; both)
- Publication-ready description of the reconstruction technique (free-text field, 500 characters limit)
- Publications that describe the algorithm
- Algorithm publicly available? (free-text field to provide the link to the website)
- If your algorithm is not yet publicly available, would you be willing to make it available at the end of the challenge? (yes, no, maybe, is already publicly available).
- Specific information about this solution (free-text field to provide specific parametric settings used)

In addition, participants were required to grant permission to the committee to publish uploaded files after the completion of the challenge.

### Submission of solutions for Stage 2

Participation in Stage 2 was restricted to participants of Stage 1. We informed participants on August 3rd via our email distribution list where data for Stage 2 may be downloaded from. Submission of solutions was performed through a dedicated online form (Google Forms), which provided detailed instructions on how to name uploaded archive files and requested the following mandatory information:

- E-mail address
- Submission Identifier of the corresponding Stage 1 submission
- File name of the zip-file you are going to upload (free-text field)
- Changes with respect to Stage 1 submission (free-text field to describe how submission differs from corresponding submission in Stage 1)
- Confirmation that the algorithm does not incorporate explicit knowledge of the ground truth and the ground was used only to optimize/adjust general algorithmic parameters.

Participants were required to grant permission to the committee to publish uploaded files after the completion of the challenge.

### Participation Statistics

We used a link management platform (Bitly) to track downloads.

### Metrics: Extended details

The metrics chosen for evaluation were:

- NRMSE: Normalized RMSE, inside the ROI. Normalization is performed by the L2-norm of the respective ground-truth.

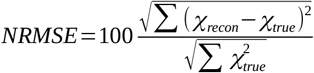
- dNRMSE: Some algorithms are known to produce underestimated results. To address this issue, we included a data demeaned and detrended RMSE score.

Demeaning and detrending was performed using the “polyfit” and “polyval” functions in MATLAB:

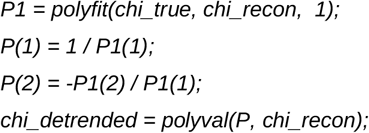

Then,

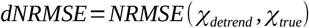

- dNRMSE Tissue: dnRMSE specific to White Matter and Grey Matter tissues.
- dNRMSE DeepGM: dnRMSE specific to Deep Grey Matter structures.
- dNRMSE Blood: dnRMSE for blood regions (effectively it was a dilated version of the vein mask).

For these three metrics, specific M_ROI_ binary masks were generated. Then:

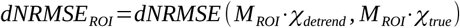

- Deviation From Linear Slope: Absolute error of the slope, derived from the demeaning and detrending process (see above). Note that this linear slope is calculated for a ROI based on the Deep Gray Matter.

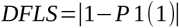
- Calcification Streak: Error metric based on the standard deviation inside neighborhood surrounding the calcification, in the difference map. To create this neighborhood, first a square mask (SQUARE) is created around the calcification. This is dilated 3 voxels in each direction. This dilated volume is then dilated other 4 voxels. The volume defined by the last dilation is assigned to a RIM mask. Then, the Calcification Streak metric is calculated by:

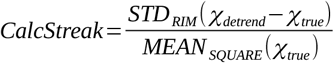
- Deviation From Calcification Moment (CalcificationError): Error in the quantification of the total moment of the calcification, defined as the volume of the reconstructed calcification multiplied by its mean susceptibility

The calcification is segmented by thresholding the reconstruction data inside the region previously defined by SQUARE. The threshold is defined as the maximum susceptibily value that contains only information inside SQUARE (does not contain information outside), starting at −1.5 ppm with 0.01ppm steps. This segmentation (CALC binary mask) is used to estimate the calcification moment by:

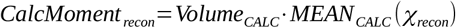

Finally,

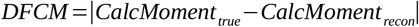

### Visual Rating GUI

**Supporting Information Figure S1.**
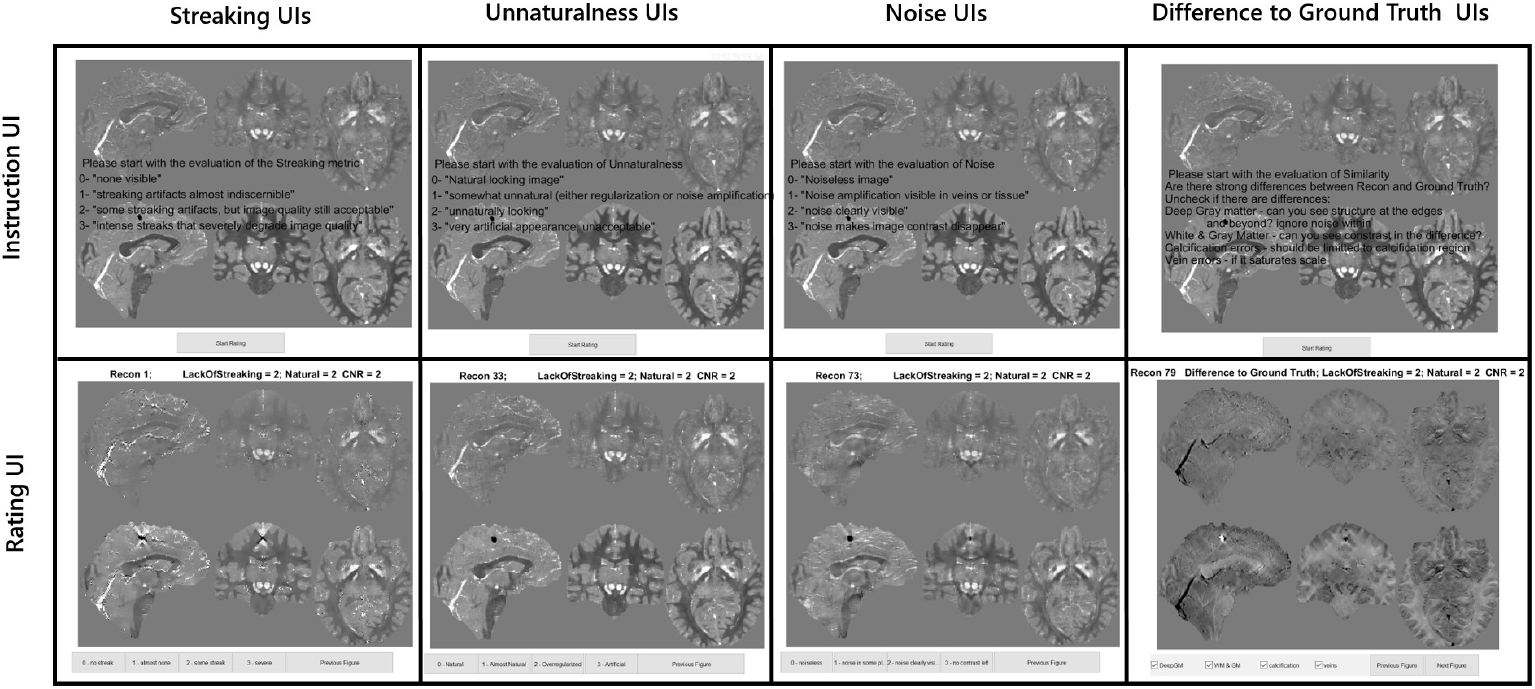
This shows an example of the simple MAtlab GUIs developed for an efficient rating process. Each rater rated one category at a time, while going through all the submissions 103 in the challenge (+ ground truth). The instruction GUI, top row appeared at the start of the rating process and remained in background in case of need to remind the criteria of rating. The Rating GUI showed three orthogonal slices of the two phantoms (SIM1 and SIM2). Raters did not have access to other slices of the reconstruction and the slices were chosen because they allowed a clear visualisation of the vein in both the sagittal plane and transverse plane, the calcification in the sagittal and coronal plane and deep grey matter in the coronal and transverse planes. For the first three classes, a mouse press on the desired rating would bring the rater to the next submission. It was possible to go back to the previous rated figure to re-rate in case of mistake, in which case the current attribute ranking to that figure would appear at the top of the GUI.

### Visual scores description

Scores from 0 (best) to 3 (worst) were given depending on the artifact level in three distinct categories, which are described in detail below.

- **Streaking** (To what extent is the submission degraded by streaking artifacts?)

- 0 - none visible
- 1 - streaking artifacts almost indiscernible
- 2 - some streaking artifacts, but image quality still acceptable
- 3 - intense streaks that severely degrade image quality
- **Unnaturalness** (How much is submission affected by features that would not appear in standard medical images? Examples for unnatural degradations include image modulation in regions that should be constant, overregularization leading to patchy, comic-like appearance, strong variations of noise-level, excessive blurring, etc.)

- 0 - Natural looking image
- 1 - somewhat unnatural, but still acceptable
- 2 - unnaturally looking
- 3 - very artificial appearance, unacceptable
- **Noise** (To what extent does noise affect the image?)

- 0 - Noiseless
- 1 - Some noise amplification visible in veins or tissue
- 2 - noise clearly visible
- 3 - noise makes image contrast disappear

For the Visual Discrepancy metrics, the raters had to identify which of the regions were clearly visible on the difference image:

- WMGM: Contrast between cortical Gray Matter and White Matter.
- DGM: Deep Gray Matter regions or structures.
- Veins: Vessels and Tissue Surrounding the Calcification (in Sim2 only).

Images that were identified to contain such information were scored “1”, being “0” otherwise, for each category.

## Supporting Results

### Participation and Submission Statistics

### Downloads and participating countries

The stage 1 data was downloaded 253 times (62 Korea, 55 USA, 39 China, 97 other countries) and we received 98 unique submissions (excluding 30 re-submissions). Of the submissions that entered the analysis phase, most were received from institutions in the US (34%), Chile (15%), Canada (12%), China (10%), and Korea (11%). Remaining submissions were from institutions in the UK (7%), Australia (3%), Germany (4%), Austria (2%), and France (1%).

The stage 2 data was downloaded 101 times (45 USA, 14 Korea, 9 Germany, 33 other countries) and we received 47 unique submissions (excluding 67 re-submissions). Of the submissions that entered the analysis phase, most submissions were received from institutions in Chile (26%), United States (23%), United Kingdom (19%), and Korea (13%). The remaining submissions were from institutions in Australia (6%), China (4%), Germany (4%), Austria (2%), and France (2%).

### Algorithms

In Stage 1, the self-reported algorithm type was “spatial-domain iterative reconstruction” for 43% of submissions and “deep learning” for 41%. The remaining submissions were “closed-form direct solutions” (5.1%), belonged to multiple categories (“Hybrid”; 4.8%), used “Inverse filtering (TKD-like)” (3.0%). 4% of submissions specified other categories. 58% of the submissions reported that the algorithm incorporated magnitude information. 54% of the submissions relied on the provided individual echo phase images, 33% relied on the provided frequency map, and 13% used both.

15% of the submissions used only a (Generalized) Total Variation regularization term, 9% only L1, and 8% only L2 regularization terms. 15% of the submissions used Total Variation in combination with other norms. 4% of the submissions used a combination of L1 and L2 terms. 50% of the solutions either did not specify the regularization approach, used implicit regularization (e.g. Deep Learning), or used more exotic approaches.

In Stage 2, the self-reported algorithm type was “spatial-domain iterative reconstruction” for 45%, similar to Stage 1. The number of submissions that used “deep learning” declined to 36%. The remaining submissions were “closed-form direct solutions” (11%) or belonged to multiple categories (“Hybrid”; 4.3%). 4% of submissions specified other categories. Similar to Stage 1, 47% of the submissions reported that the algorithm incorporated magnitude information. The percentage of submissions that relied on the provided individual echo phase images increased to 68%, 32% relied on the provided frequency map, and none used both.

**Supporting Information Table S1.**
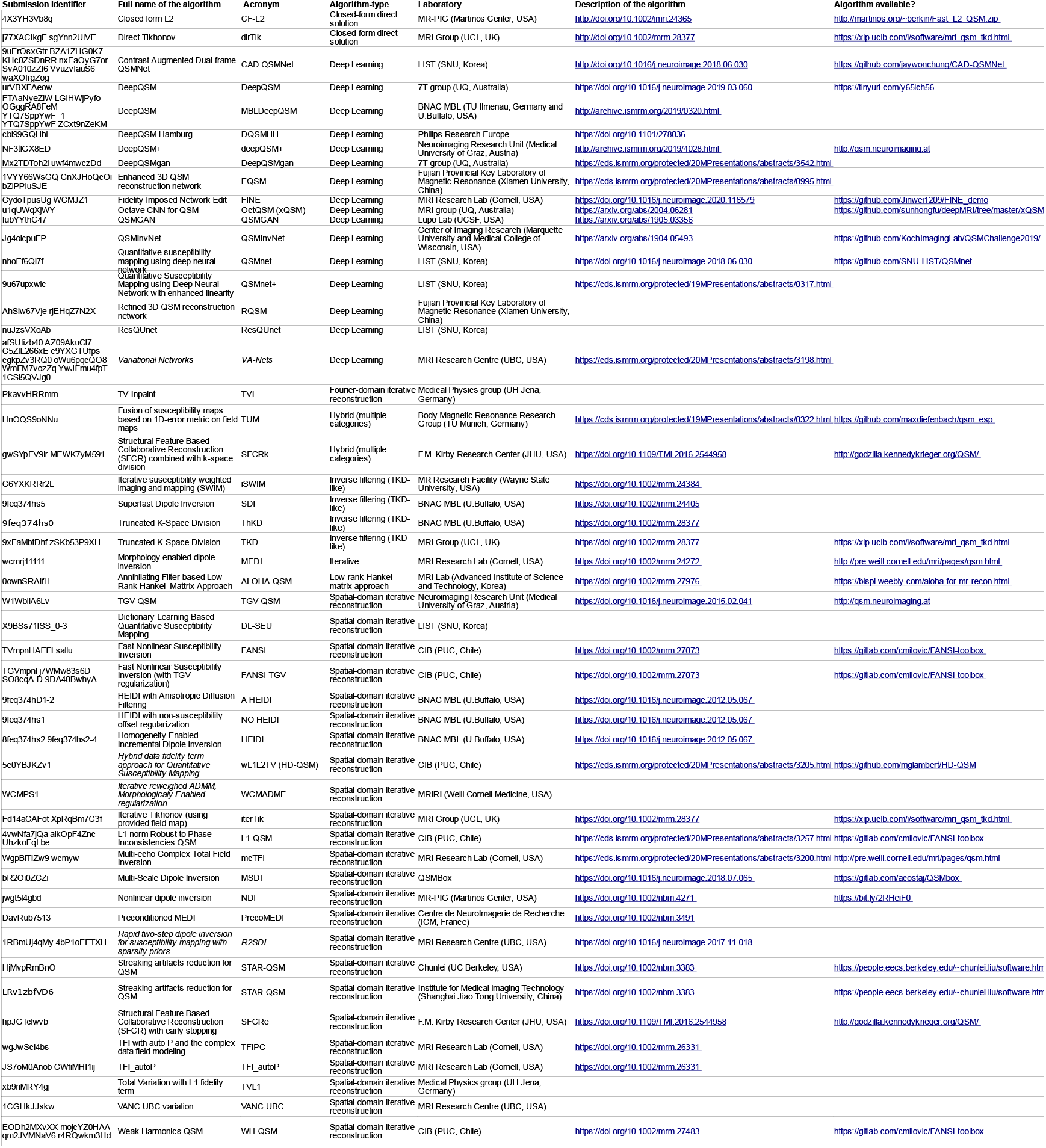
Publications that describe each submitted algorithm, and code repositories, if available. Multiple submissions with the same algorithm (but significantly different parameters or results) are grouped.Results of Stage 1

**Supporting Information Figure S2.**
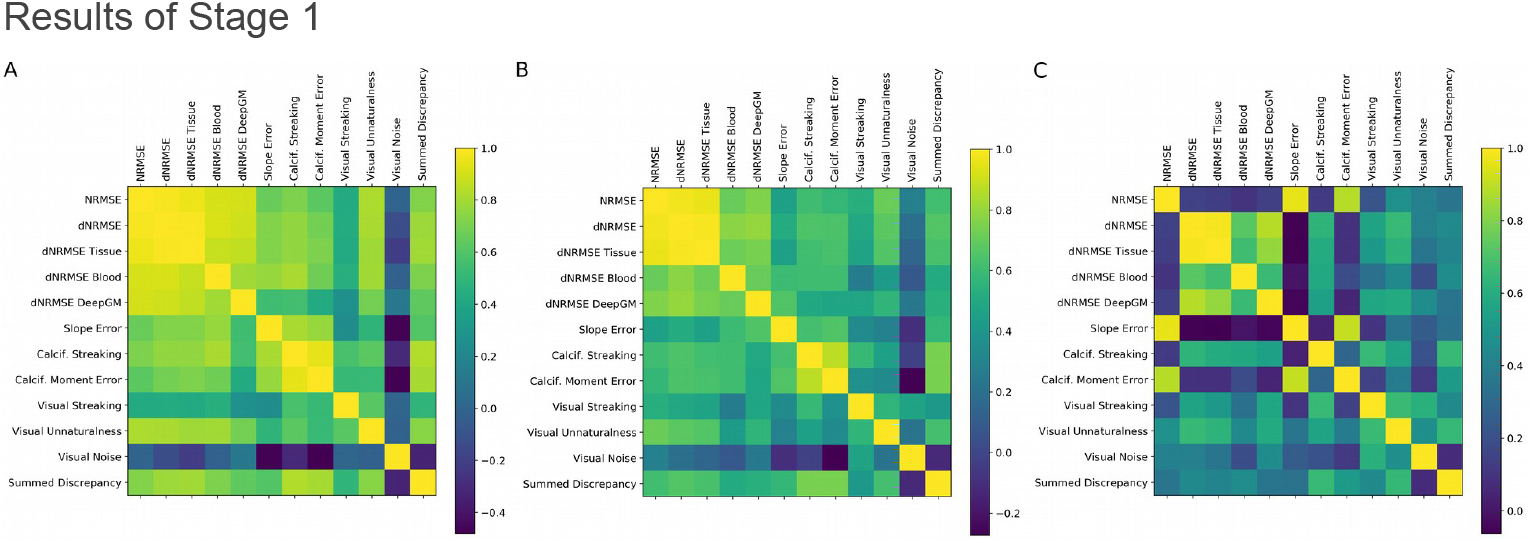
Stage 1 correlation between official metrics and visual assesment for A) top 5 submissions, B) submissions with NMRSE<80, and C) all submissions.

**Supporting Information Figure S3.**
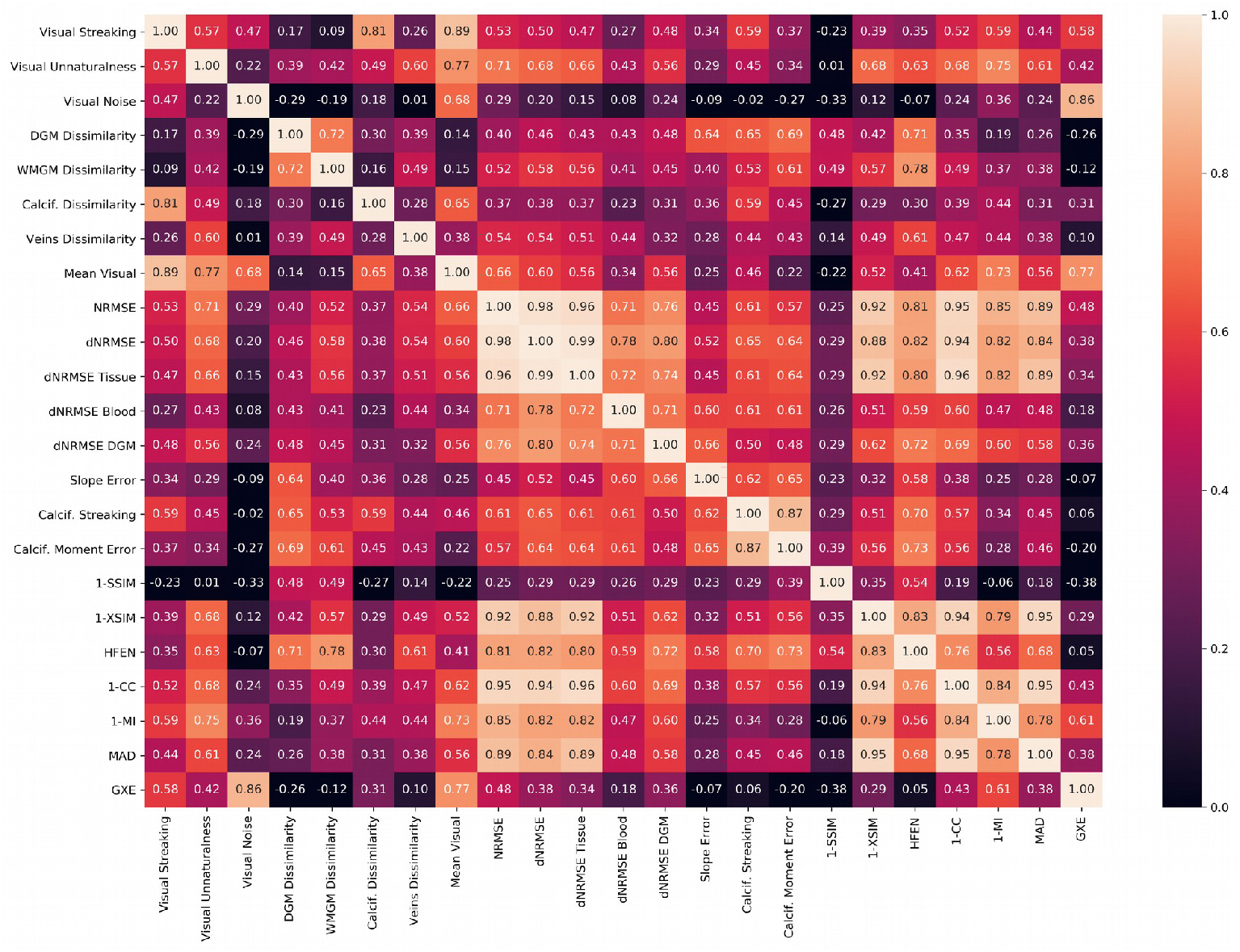
Correlation between official, visual and extended global metrics, for all submissions with NMRSE<80 in Stage 1.

**Supporting Information Figure S4.**
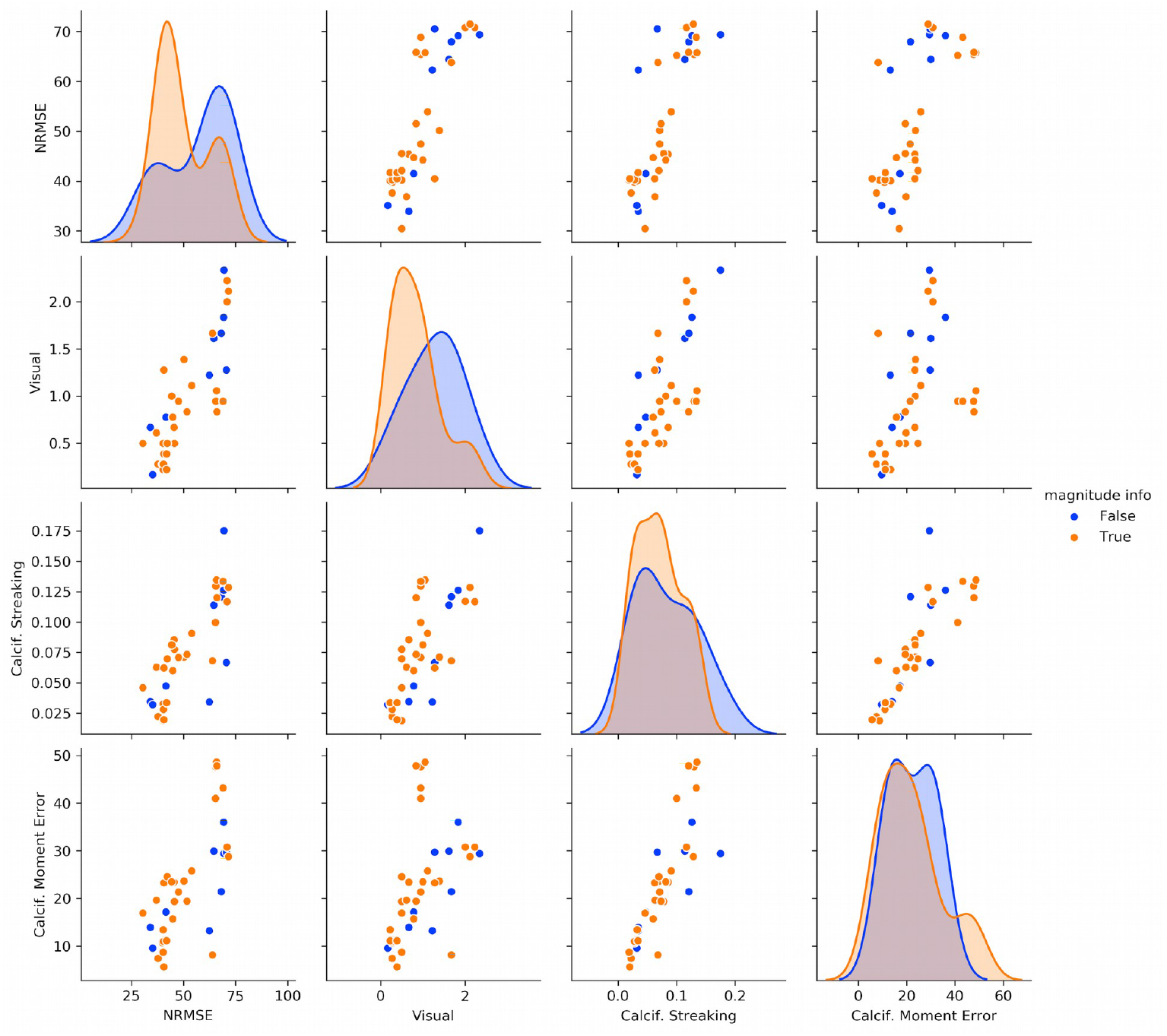
Stage 1 scatterplots between selected pairs of metrics comparing the use of magnitude information as prior or weighting for the reconstruction. The diagonal shows estimated histograms for each metric.

**Supporting Information Figure S5.**
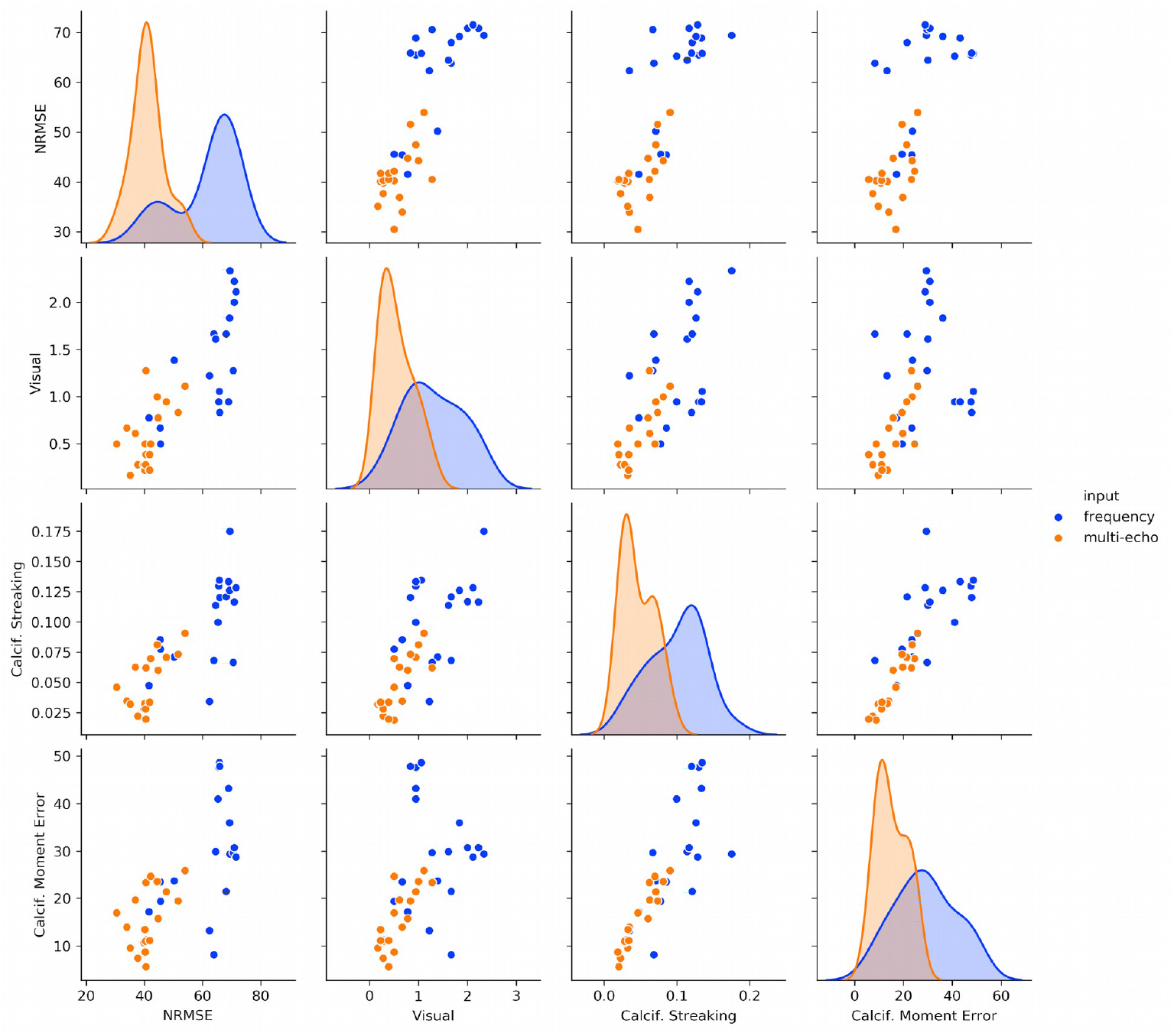
Stage 1 scatterplots between selected pairs of metrics comparing the use of multi-echo and frequency maps as input for reconstruction. The diagonal shows estimated histograms for each metric. Overall, algorithms using multiecho data showed lower errors.

**Supporting Information Figure S6.**
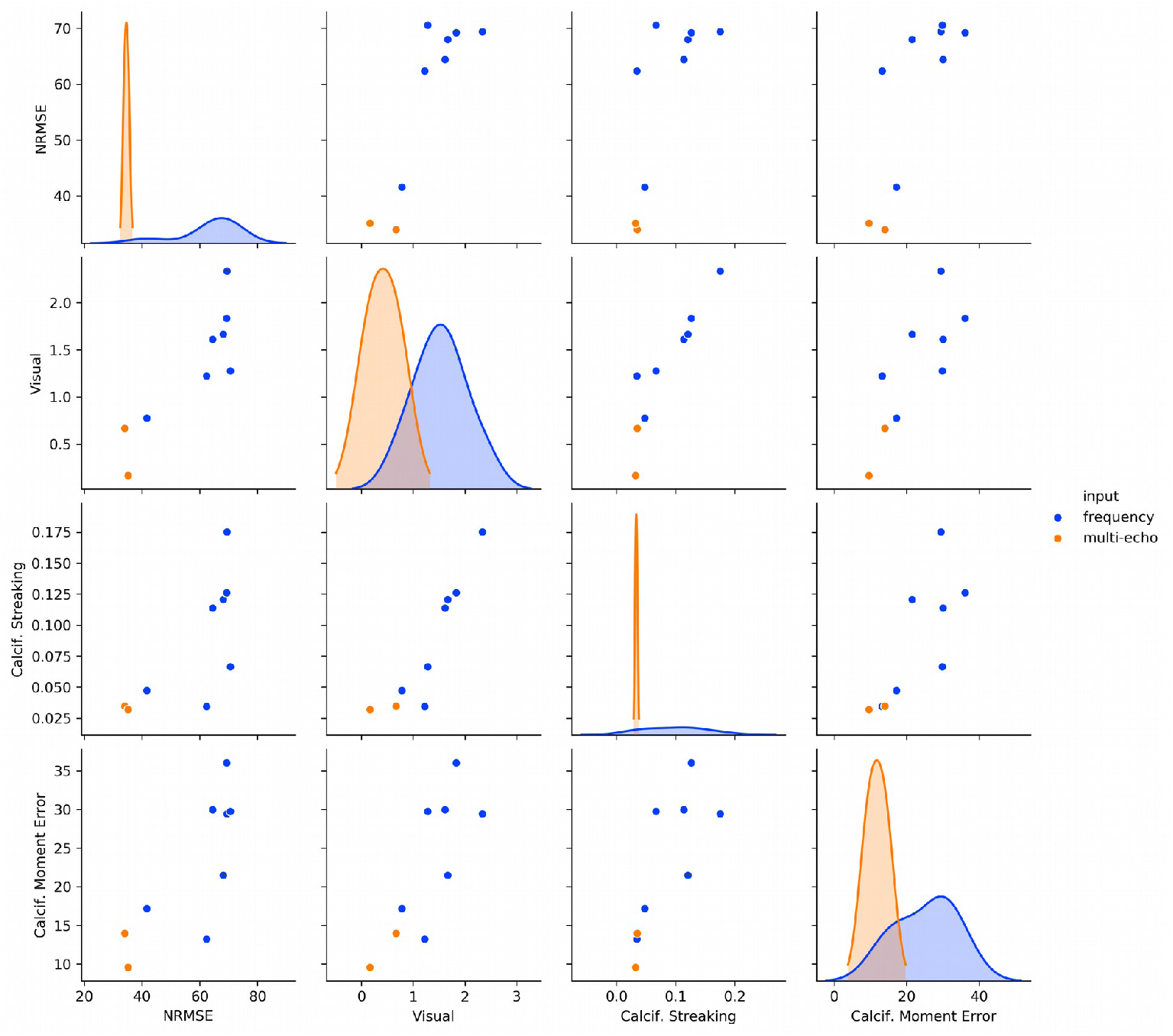
Stage 1 scatterplots between selected pairs of metrics comparing the use of multi-echo and frequency maps as input for reconstruction for top 9 submissions that didn’t include the magnitude information as prior or weight.

**Supporting Information Figure S7.**
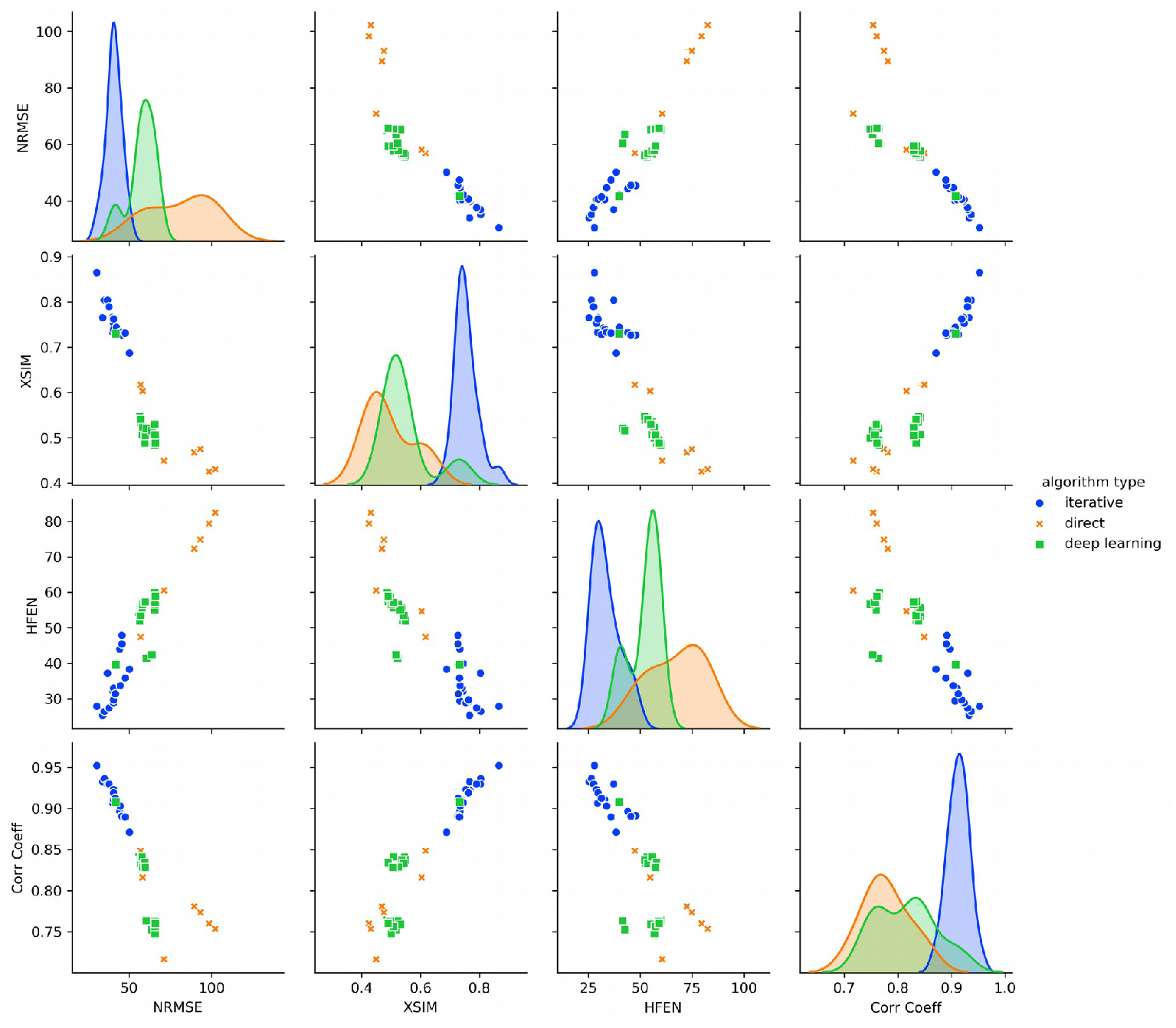
Stage 1 scatterplots between NMRE and extended metrics XSIM, HFEN and Correlation Coefficient. Only the best 20 RMSE submissions for each algorithm type are shown.

**Supporting Information Figure S8.**
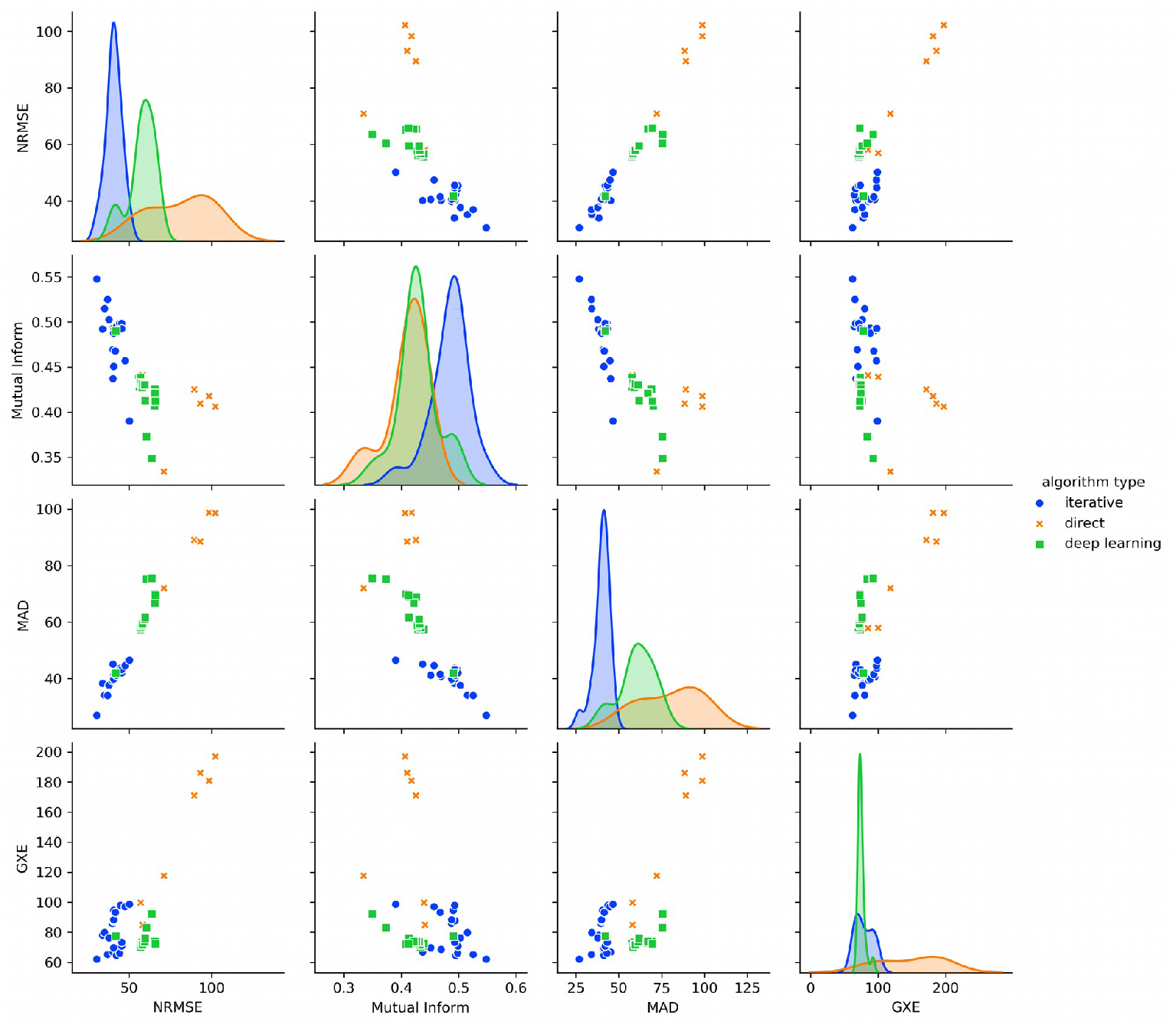
Stage 1 scatterplots between NMRE and extended metrics Mutual Information, Mean Absolute Error (MAD) and RMSE measured on the gradient (first derivative) domain (GXE). Only the best 20 RMSE submissions for each algorithm type are shown.

**Supporting Information Table S2.**
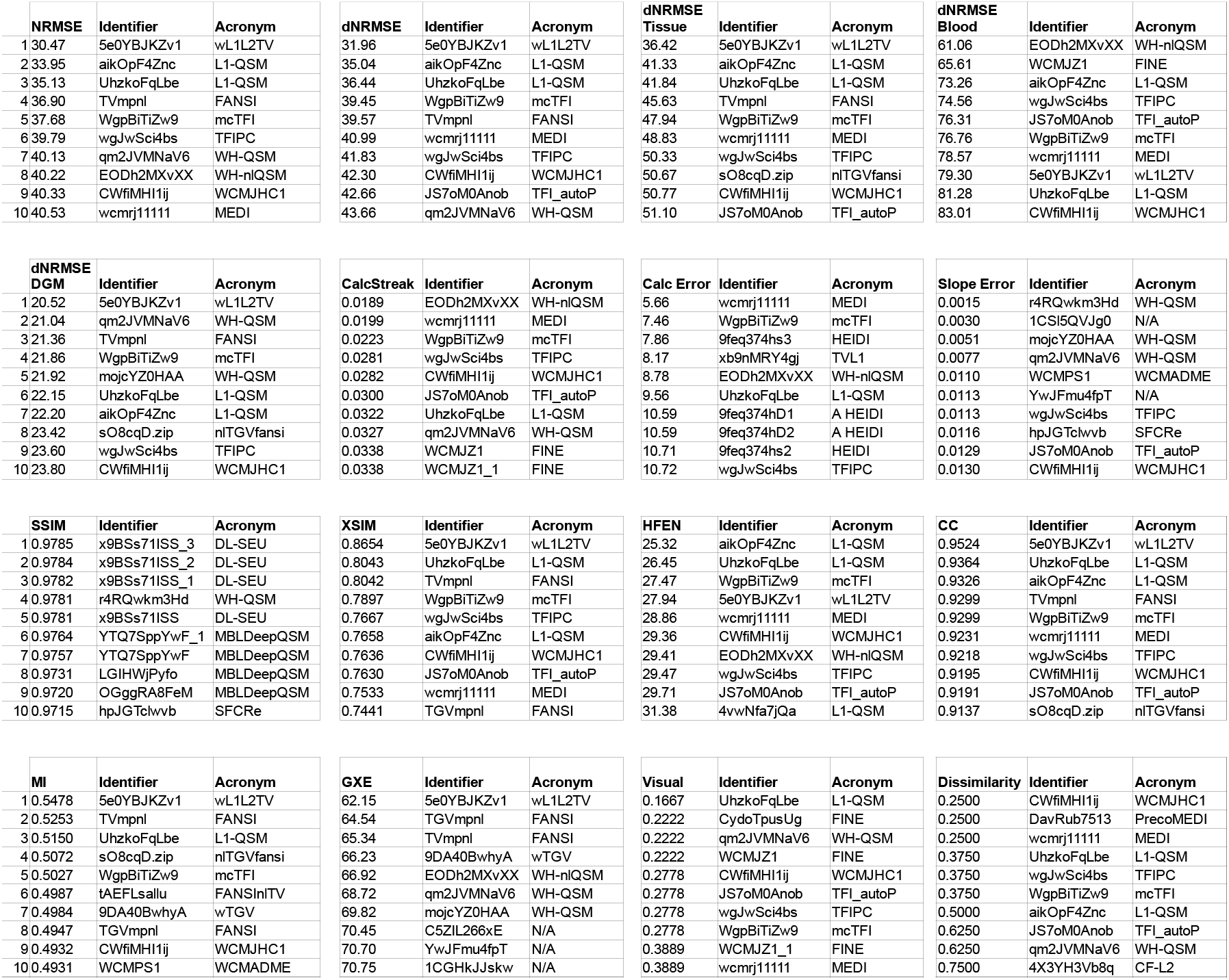
Top 10 best scoring submissions and scores in Stage 1, for each measured metric (official and extended).

**Information Figure S9.**
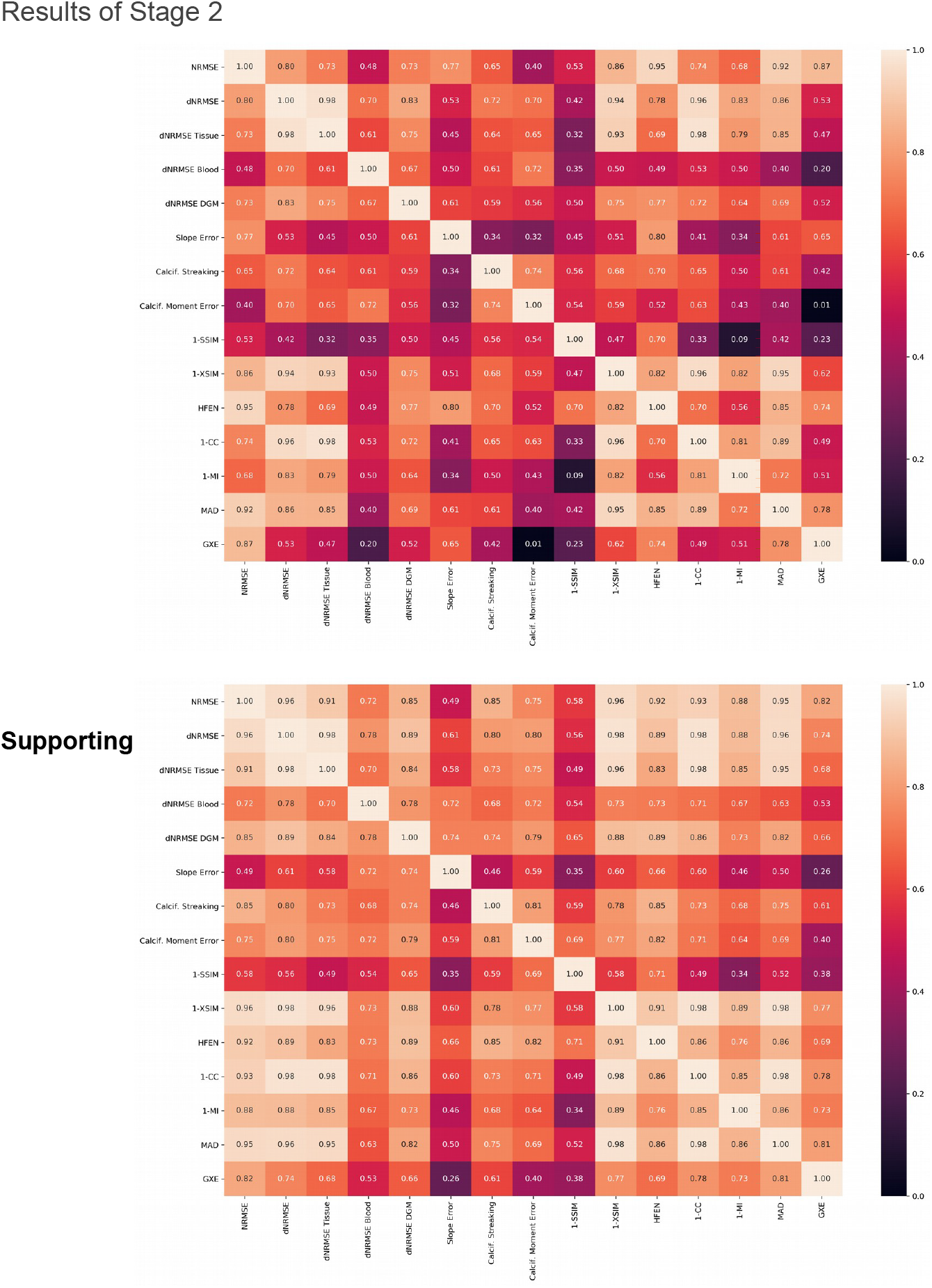
Correlation matrix for Stage 1 submissions that corresponds to submissions to Stage 2 (top) and all Stage 2 submissions (bottom). Correlations between metrics significantly increased in comparison to Stage 1.

**Supporting Information Figure S10.**
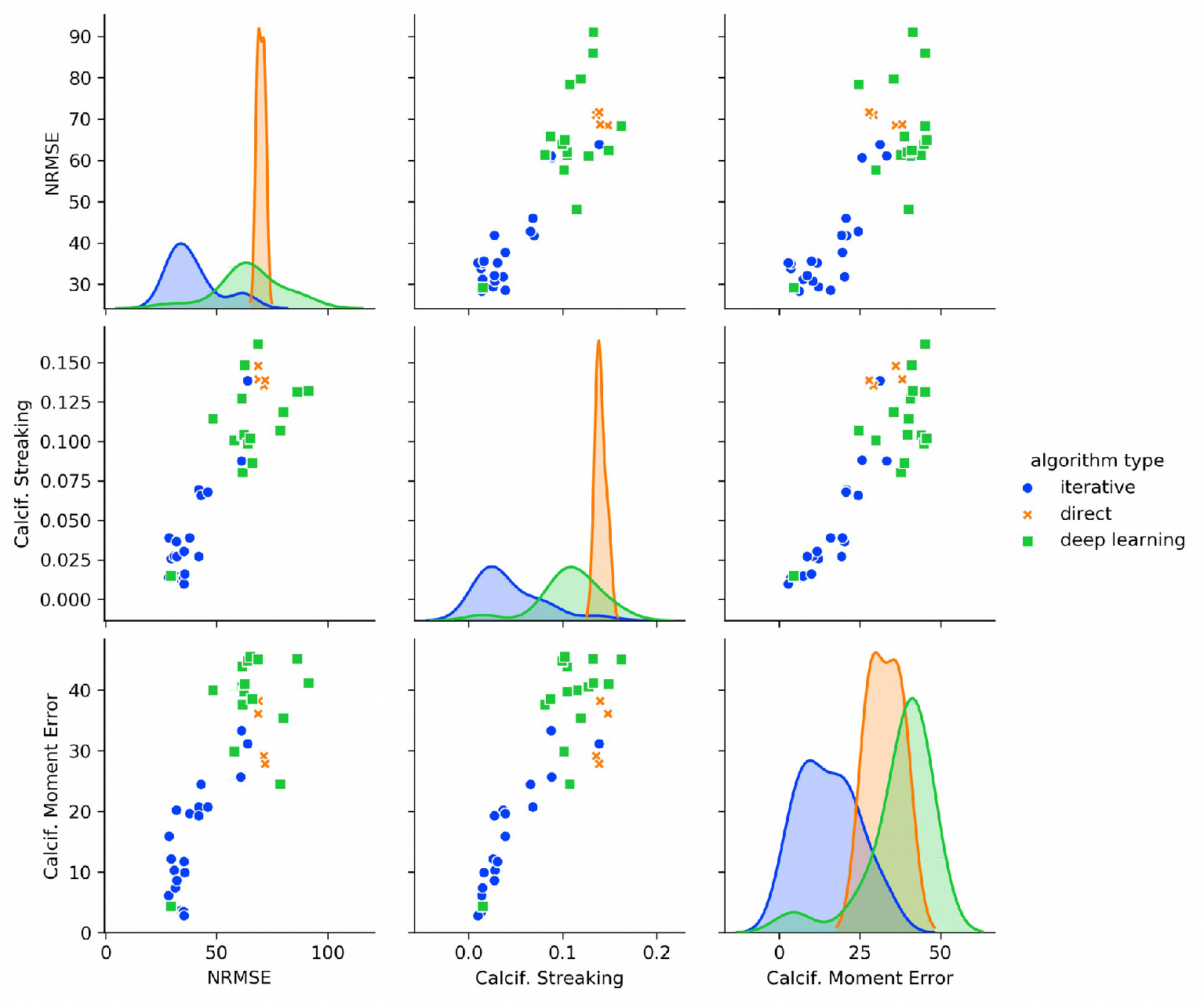
Stage 2 scatterplots between selected pairs of metrics showing the top 20 submissions in each algorithm class (shown as different colors, see legend). The diagonal shows estimated histograms for each metric. Overall, iterative methods significantly performed better than Deep Learning and Direct approaches, showing larger relative improvements (see main Figure 5).

**Supporting Information Figure S11.**
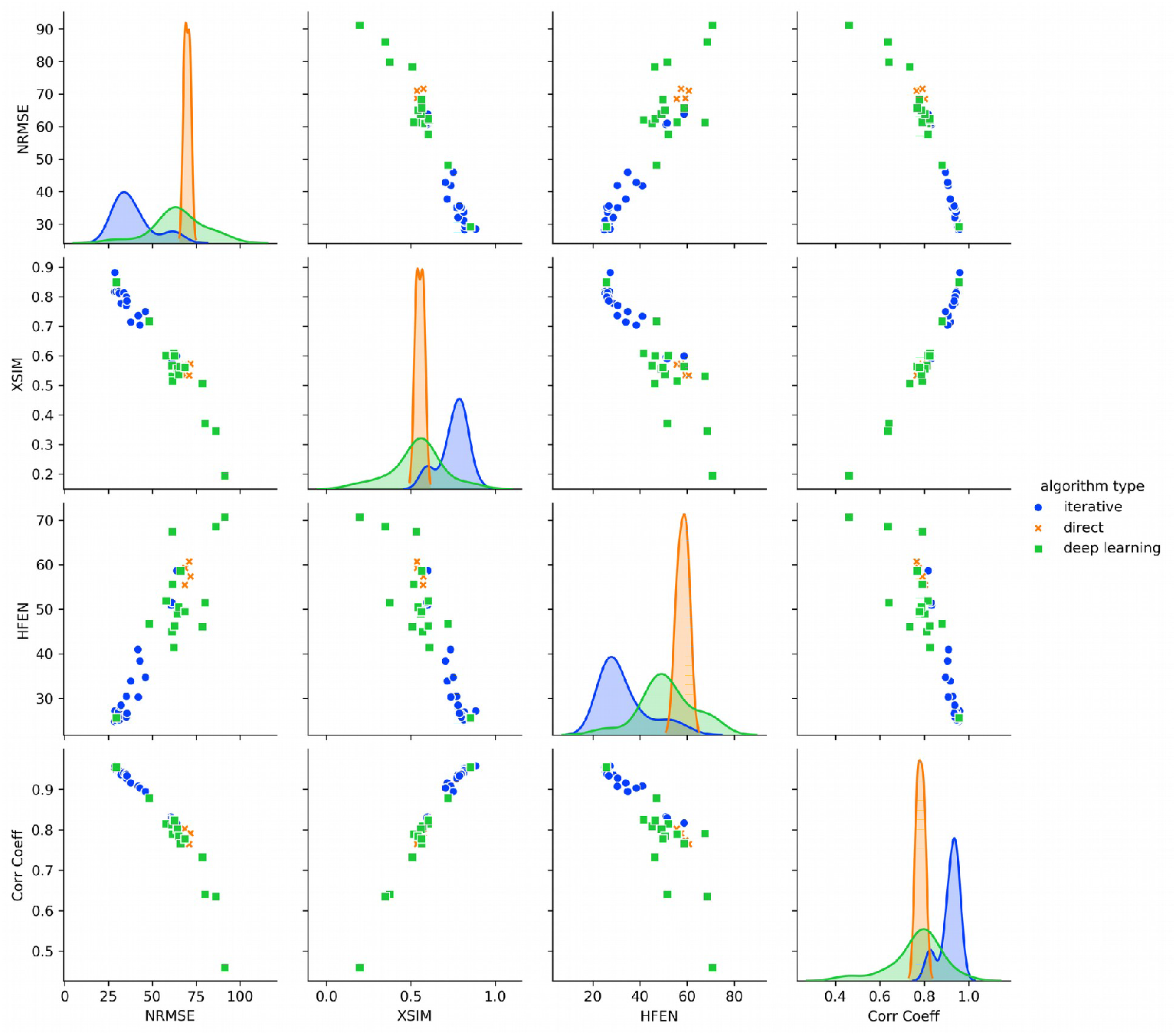
Stage 2 scatterplots between NMRE and extended metrics XSIM, HFEN and Correlation Coefficient. Only the best 20 RMSE submissions for each algorithm type are shown.

**Supporting Information Figure S12.**
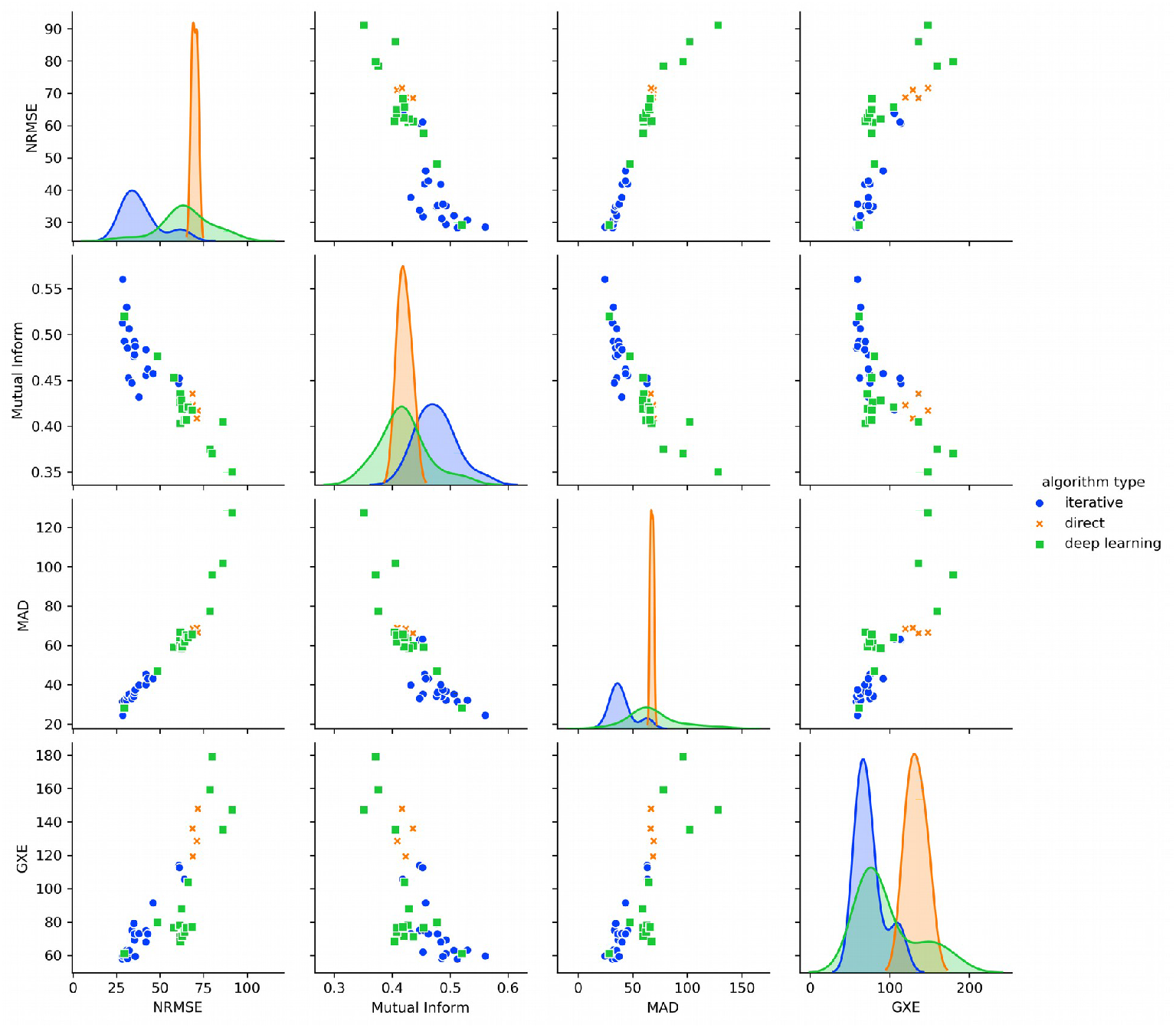
Stage 1 scatterplots between NMRE and extended metrics Mutual Information, Mean Absolute Error (MAD) and RMSE measured on the gradient (first derivative) domain (GXE). Only the best 20 RMSE submissions for each algorithm type are shown.

**Supporting Information Table S3.**
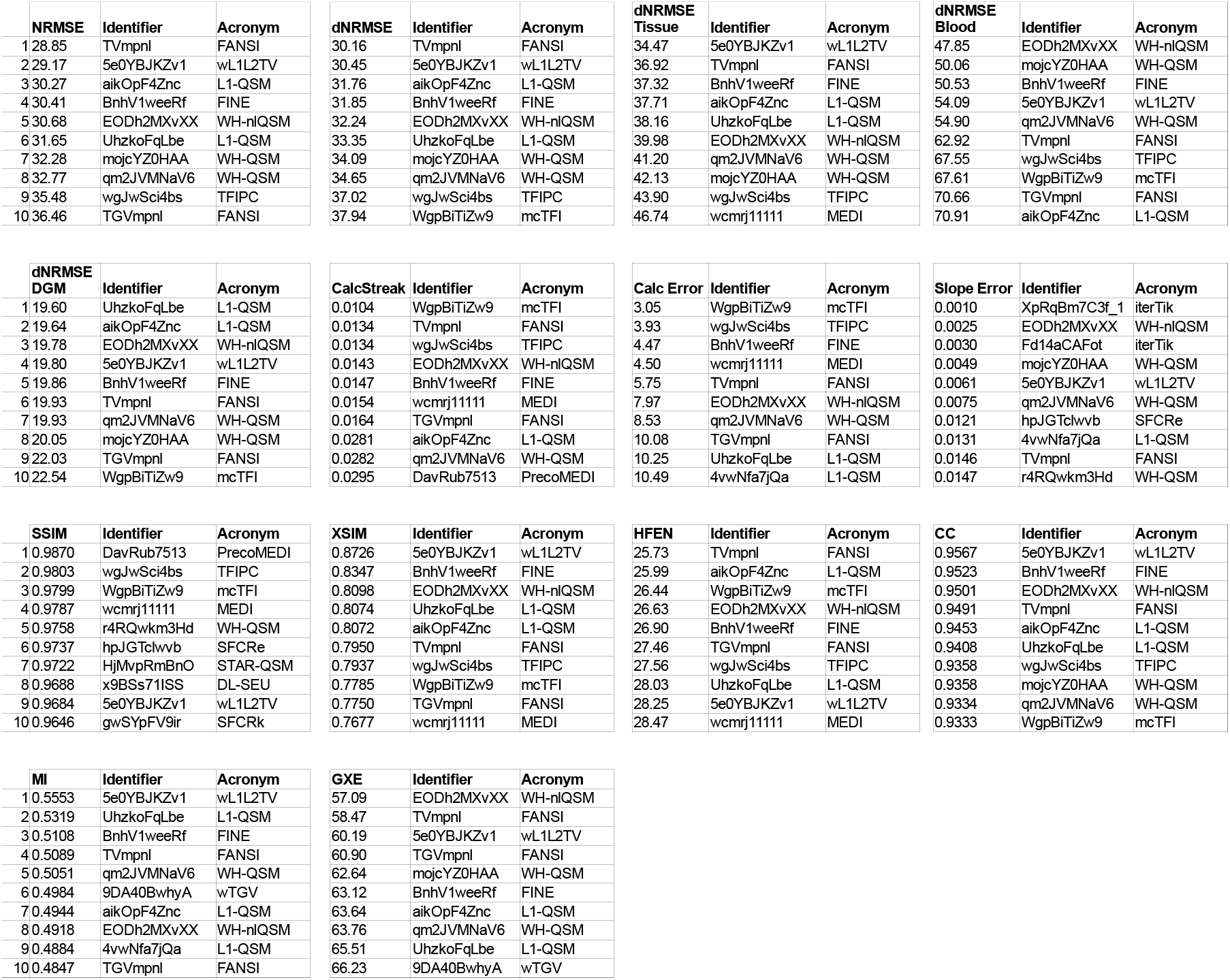
Top 10 best scoring submissions and scores in Stage 2, for each measured metric (official and extended).

